# A model for cortical activity sequences

**DOI:** 10.1101/2024.02.25.581959

**Authors:** Andrew B. Lehr, Finn Erzmann, Carlo Michaelis, Julia Nowak, Alexander Gail, Arvind Kumar, Christian Tetzlaff

**Affiliations:** Department of Computational Neuroscience, University of Göttingen, Germany; Group of Computational Synaptic Physiology, Department of Neuro- and Sensory Physiology, University Medical Center Göttingen, Germany; Sensorimotor Group, German Primate Center, Göttingen, Germany; Department of Computational Science and Technology, KTH Royal Institute of Technology, Sweden

## Abstract

Networks of neurons in the brain, that act on a timescale of milliseconds, can intrinsically generate reliable sequential activity on slow behavioral timescales of seconds. A possible mechanism for intrinsic sequence generation based on theoretical evidence points to distance-dependent connectivity with correlated spatial asymmetries, establishing an anisotropic network connectivity. We show that networks with such correlated asymmetric connectivity as well as symmetric distance-dependent connectivity match experimental data of connectivity motifs as well as neuronal activity statistics from rat and monkey cortex. At the network level, however, only the correlated asymmetric connectivity pattern generates spatiotemporal activity sequences on behaviorally relevant timescales, while the symmetric connectivity results in transient but stationary spatial bumps of neural activity. Our results strongly support the role of correlated asymmetries in connectivity for the generation of sequential activity in neural networks.

## 1 Introduction

Spatiotemporal sequences of neuronal activation have emerged as a key feature of network dynamics in the brain. Sequential activity has been observed across brain regions and animal species in various behavioral tasks such as decision making^1,2^, timing^3,4^, olfactory processing^5^, birdsong generation^6^, motor control^7–9^, and hippocampal-dependent learning and memory^10–19^. Reproducible sequential activity can span hundreds of milliseconds up to minutes^20^, orders of magnitude longer than the timescale of single spikes. Crucially, sequences arise even without sequential input^1,2,6,11,17,21^ suggesting that they can be intrinsically generated by the local network structure. However, the theoretical basis of how recurrently and locally connected spiking neurons can generate such sequences is a longstanding matter of active investigation^22–41^.

Experimental observations have revealed features of cortical connectivity thought to support intrinsic spatiotemporal sequence generation. The probability of a connection between neurons is distance-dependent, decreasing on a scale of hundreds of micrometers^42–46^. Therefore, a neuron has a higher probability to connect to its close neighbours. Neurons can furthermore have asymmetric projection patterns with axons or dendrites extending preferentially in a particular direction^43–48^. In mouse visual cortex, pyramidal neurons receive spatially asymmetric inputs that determine directional tuning^46,49^, with nearby neurons that were born together sharing inputs^50^. In mouse sensory cortex nearby neurons project preferentially in similar directions aligned with the propagation direction of travelling waves^51^, spatiotemporal sequences of neuronal activation that have been observed in many settings^52–71^.

From locally connected recurrent network models (LCRN), it is well known that symmetric distance-dependent connectivity supports the formation of spatially localized neuronal activity, referred to as “bumps”^22,28,31,36,37,72,73^. Furthermore, Spreizer et al.^37^ show that a rather small, but correlated asymmetry in the distance-dependent connectivity can induce a movement of the localized bumps through the network resulting in spatiotemporal sequences on a timescale of several seconds. Based on these results, here, we perform a more detailed comparison to microcircuit connectivity data and neuronal firing statistics to demonstrate that correlated asymmetric connectivity provides a biologically reasonable model underlying the generation of neural spatiotemporal sequences.

On the microcircuits level, connections between neurons are not random but instead show enhanced connectivity motifs like reciprocal connections, chains, convergence and divergence^42,45,74–77^. We show that locally connected networks reproduce the nonrandom connectivity motifs observed in cortex^74^ regardless of whether projections are symmetric or correlated asymmetric. Further, by introducing heterogeneity in neuronal timescales^78^ via a distribution of refractory periods^79^, we show that neuronal activity resulting from both symmetric and correlated asymmetric connectivity can reliably match monkey motor/premotor cortex neuronal firing statistics. Thus, both types of connectivity, symmetric and correlated asymmetric, are consistent with cortical connectivity motifs as well as single neuron activity statistics of the cortex. At the network level, however, only correlated asymmetric connectivity generates spatiotemporal activity sequences on timescales of seconds.

## 2 Results

### 2.1 Both symmetric and asymmetric distance-dependent projections match cortical connectivity motifs

We first compared recurrent networks with distance-dependent connectivity to experimentally observed connectivity motifs from rat visual cortex^74^. We considered an *isotropic* model with symmetric distance-dependent connectivity in which a Gaussian kernel centered at each neuron defines the probability of forming a synapse with surrounding neurons. Furthermore, we considered an *anisotropic* model in which the center of the Gaussian kernel is slightly shifted introducing on average an asymmetry in the connectivity. For nearby neurons the shift and thus the asymmetry is correlated (Fig. 1a). Note that the level of asymmetry is small compared to the isotropic network (Fig. 1c), but sufficient to generate local bumps of increased activity moving through the network to form a spatiotemporal sequence (Fig. 1d). We varied two parameters in both models (Fig. 2b): (i) the connection locality, i.e. standard deviation *σ*_loc,exc_ of the Gaussian kernel for excitatory neurons, with a lower standard deviation implying more local connections, and (ii) the proportion of synapses formed randomly *p*_con,rand_, independent of the Gaussian kernel. Connection locality *σ*_loc,exc_ ranged from 0.01 to 0.25 in steps of 0.01 and the proportion of random connections *p*_con,rand_ ranged from no random connections to 25% of the overall synapses chosen randomly in steps of 1%.

**Figure 1.**
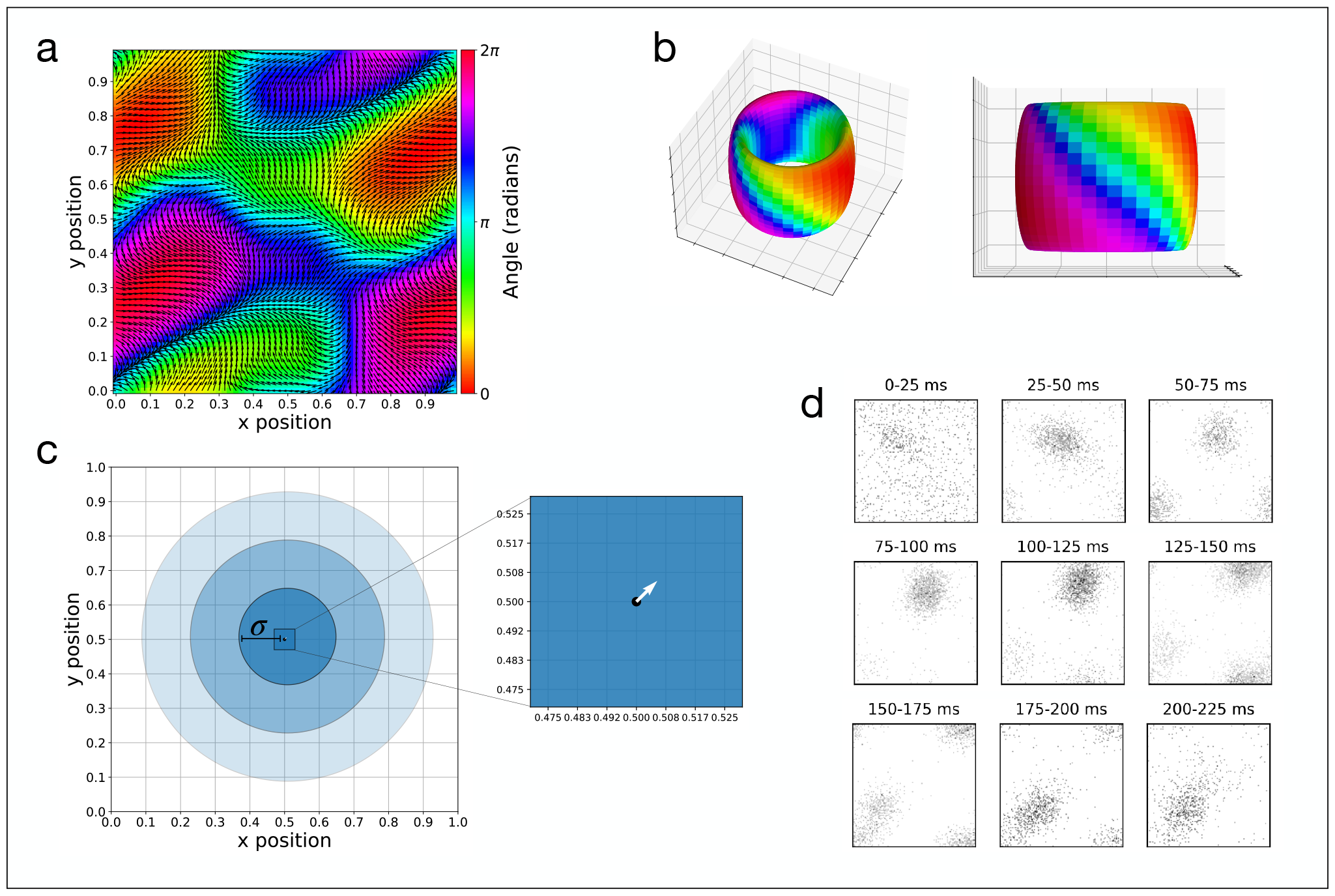
The anisotropic network model. **(a)** The Perlin connectivity landscape with preferred directions (angles) color coded and shown as a vector field. **(b)** The network has periodic boundary conditions and is thus folded to give a toroidal network topology. **(c)** An example Gaussian connectivity kernel with standard deviation *σ* = 0.14 and shift *d* = 1 is shown. Each ring is one standard deviation. Neurons are equidistantly spaced on the grid such that for a network with *N* = 14400 excitatory neurons, one of the grid squares (0.1x0.1) contains 144 excitatory neurons. The zoom shows how small the shift is in comparison to the network size. In the zoomed region, one excitatory neuron is located at each grid corner. **(d)** Under these circumstances, spatially localized activity emerges in the network and given the correlated asymmetry, becomes “unstuck in space”, moving around the network. Each frame shows all spikes from a 25 ms time window in response to a transient spatially local burst of input arriving at 10 ms. A black dot was plotted for each spike at the location the corresponding neuron.

**Figure 2.**
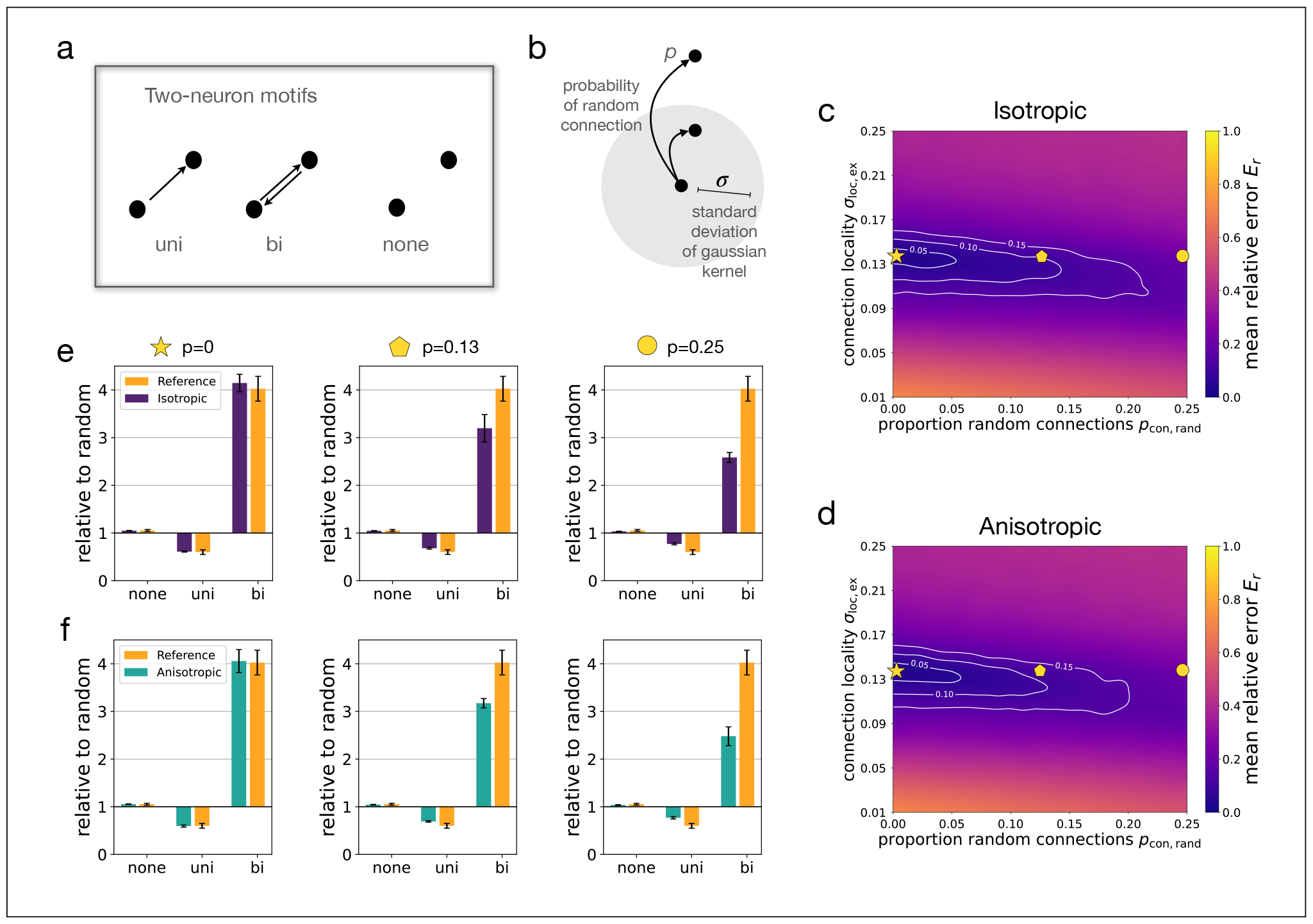
Locally connected networks fit experimentally observed two-neuron connectivity motifs. **(a)** The three possible two-neuron motifs are shown — unidirectional, bidirectional, or not connected. **(b)** Depiction of connectivity parameters that were varied. The standard deviation of the Gaussian connectivity kernel is parameterized by *σ*_loc,exc_ and the probability of a synapse being a random connection not defined by the Gaussian kernel is *p*_con,rand_ (shortened to *σ* and *p* in figure). **(c)** Mean relative error between two neuron connectivity motifs measured in the isotropic network model vs. rat visual cortex data^74^. **(d)** Same as (c) but for the anisotropic network model. Both (c) and (d) are averaged over 5 randomly generated networks drawn according to the parameters *σ*_loc,exc_ and *p*_con,rand_. Graphs are smoothed and contours are added to aid visualization (see Methods). Shapes indicate parameter sets shown in (e,f). **(e)** Two-neuron connectivity motifs relative to expectation based on random connectivity is shown for the isotropic model (purple) for three examples each with *σ*_loc,exc_ = 0.14 and increasing random connectivity *p*_con,rand_ = 0, *p*_con,rand_ = 0.13, and *p*_con,rand_ = 0.25, compared to experimental data^74^ (yellow). **(f)** Same as (e) but for the anisotropic network model (teal). Bars show mean *±* standard deviation over *N*_seed_ = 5.

#### Two-neuron motifs

For pairs of neurons there are three possible motifs: bidirectional, unidirectional or no connection (Fig. 2a). In rat visual cortex data, Song et al.^74^ observed that unidirectional connections are underrepresented while bidrectional connections are highly overrepresented compared to what is expected in a randomly connected network. We found that this result can be accurately reproduced by both the *isotropic* and *anisotropic* models (Fig. 2c and 2d, respectively). In both cases, a large region of the parameter space fit the data well (purple region and white contours, Fig. 2c,d). Both models fit the cortical motif data best for connection locality around *σ*_loc,exc_ = 0.14 and with very low or no probability of random connections *p*_con,rand_ ≈ 0 (see Fig. 2c,d). Note that correlated asymmetry in the projections had no observable effect on the motifs evidenced by the similarity of the *isotropic* and *anisotropic* results. The fit deteriorates when the proportion of random connections is increased (Fig. 2e,f), which is to be expected since for a randomly connected network with *p*_con,rand_ = 1, all over- and underrepresentations should vanish, given that the measure is relative to motif counts in a random network.

#### Three-neuron motifs

For groups of three neurons, we again estimated the mean relative error *E*_*r*_ for the same parameter space as for the two-neuron motifs (*σ*_loc,exc_ ∈ [0.01, 0.25] and *p*_con,rand_ ∈ [0, 0.25]) in both models. The underlying connection probabilities *p*_uni_, *p*_bi_ and *p*_none_, which are used to calculate the expected number of three-neuron motifs in a randomly connected network (see Methods Section 4.8 and Table 2), are chosen based on the experimental results for the two-neuron motifs from Song et al.^74^. For three-neuron motifs the region with the lowest error has a similar position within the parameter space to the results for two-neuron motifs, with anisotropic and isotropic model performing similarly (Fig. 3c,d). The values of *E*_*r*_ are however larger for three-neuron motifs indicating a less accurate match compared to the two-neuron motifs, though this is compensated by larger standard deviations of observed three-neuron motifs in the experimental data and simulation (Fig. 3e,f). Fig. 3e,f shows over- and underrepresentations of the 16 different three-neuron motifs averaged over five network initializations for each model and the experimental data for comparison. The examples are taken from the region in the parameter space with lowest error for the two-neuron motifs (purple region in Fig. 2c,d, *σ*_loc,exc_ = 0.14, *p*_con,rand_ = 0). Two- and three-neuron motif counts for the best fits on triads and pairs are shown in Supplementary Fig. S1.

**Figure 3.**
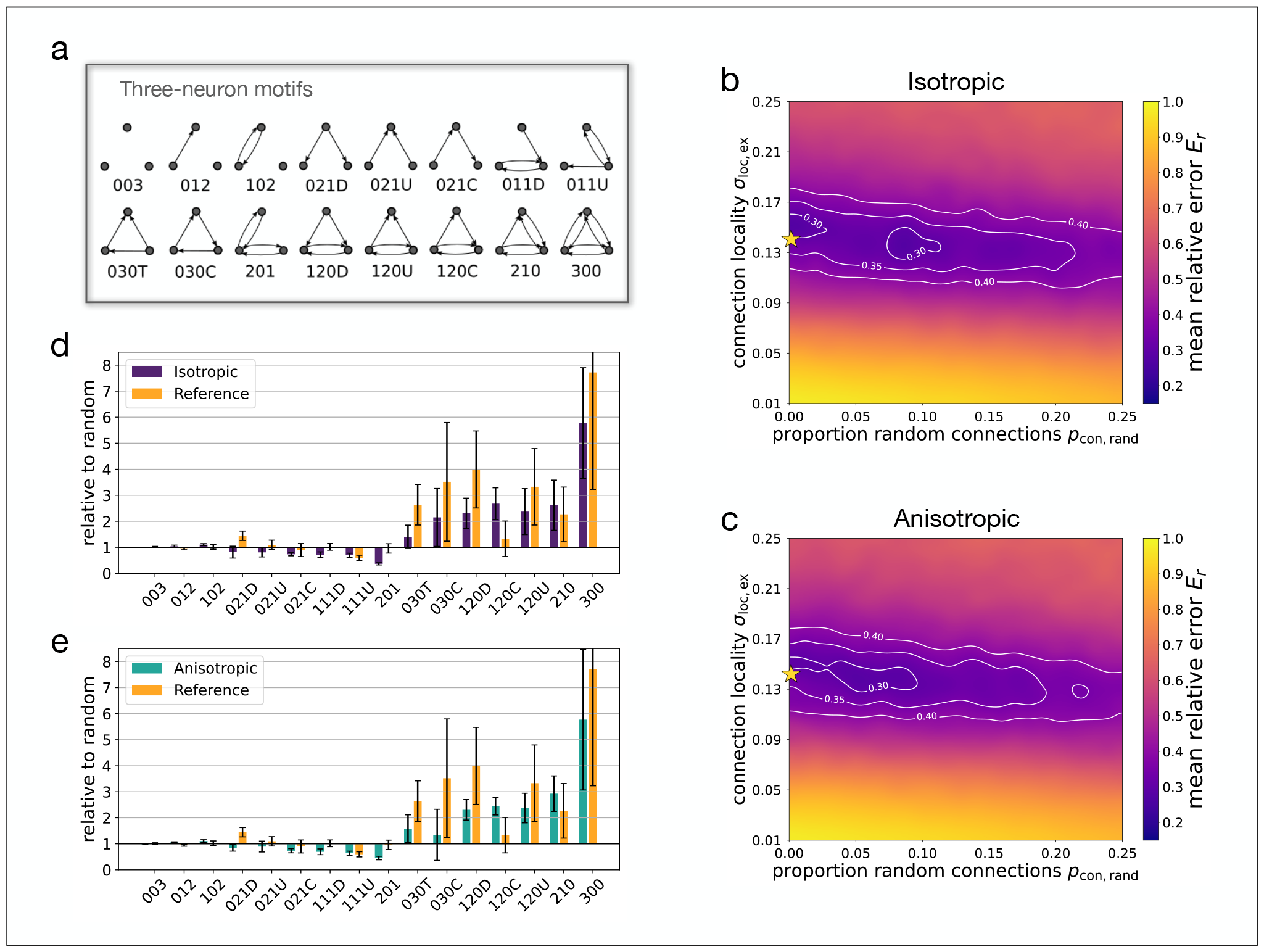
Locally connected networks fit experimentally observed three-neuron connectivity motifs. **(a)** The sixteen possible three-neuron motifs are shown. **(b)** Mean relative error between three-neuron connectivity motifs measured in the isotropic network model vs. rat visual cortex data^74^. **(c)** Same as (b) but for the anisotropic network model. Both (b) and (c) are averaged over 5 randomly generated networks drawn according to the parameters *σ*_loc,exc_ and *p*_con,rand_. Graphs are smoothed and contours are added to aid visualization (see Methods). Stars indicate parameters used for examples shown in (d,e). **(d)** Three-neuron connectivity motifs relative to expectation based on observed two-neuron motifs (see Methods and Song et al.^74^) are shown for the isotropic model (purple) for parameters identified in two-neuron motif comparison above (gold star in (b), *σ*_loc,exc_ = 0.14, *p*_con,rand_ = 0) with comparison to experimental data^74^ (yellow). **(e)** Same as (d) but for the anisotropic network model (teal). Bars show mean *±* standard deviation over *N*_seed_ = 5.

### 2.2 Neurons with distance-dependent projections and heterogeneous refractory periods match single neuron cortical activity statistics

With both the *isotropic* and *anisotropic* models being able to reproduce cortical microcircuit connectivity motifs, we next turned to neuronal activity statistics. We compared the models with experimental data from macaque monkey motor (M1) and premotor cortex (PMd) during a memory-guided center-out reach task in terms of firing rate distributions, interspike interval distributions, and the coefficient of variation (see Methods). For the simulations, connectivity in the models was based on the results of fitting the connectivity motifs, *σ*_loc,exc_ = 0.14 and *p*_con,rand_ = 0.

Since we were interested in intrinsically generated activity, we administered instantaneous spatially local bursts of input during ongoing background input. In order to fit the models to the activity statistics of experimental data, we introduced heterogeneity in neuronal timescales by considering a distribution of refractory periods *τ*_ref_, parameterized by the average refractory period *µ*_*τ*_ and the standard deviation of the refractory period distribution *σ*_*τ*_ (Fig. 4a). For each refractory period distribution (*µ*_*τ*_, *σ*_*τ*_) we ran five simulations of 500 milliseconds (based on average task duration in experimental data, see *Methods: Electrophysiological data*) for the *isotropic* and *anisotropic* networks, in which the recurrent connections and background input differed and the spatially local burst of input remained the same.

**Figure 4.**
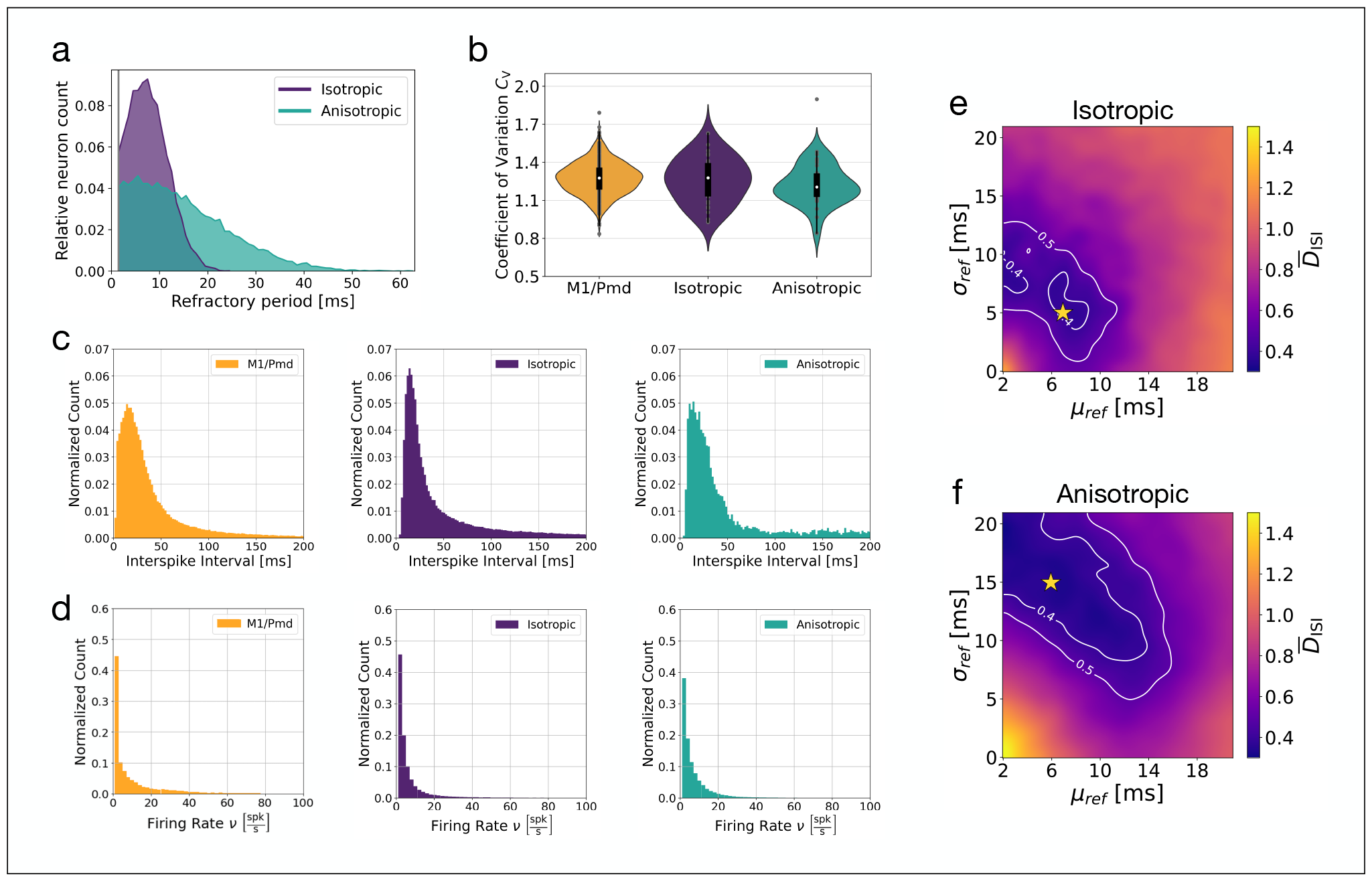
Single neuron activity statistics in isotropic and anisotropic networks with heterogeneous neurons match experimental data. **(a)** Example refractory period distributions for isotropic (*µ*_*τ*_ = 7 ms, *σ*_*τ*_ = 5 ms) and anisotropic (*µ*_*τ*_ = 6 ms, *σ*_*τ*_ = 15 ms) networks, see stars in (e) and (f). Normalized by number of excitatory neurons *n*_*e*_ = 14400. **(b)** Coefficient of variation for the M1/PMd data compared to the isotropic and anisotropic network simulations. **(c)** Interspike interval distributions for experimental data and network simulations. **(d)** Firing rate distributions for experimental data and network simulations. **(e)** Distance between the interspike interval distribution for the experimental data and isotropic network simulations as a function of the refractory period distribution (*µ*_*τ*_, *σ*_*τ*_). **(f)** Same as (e) but for the anisotropic network. Panels (a-d) correspond to stars in (e) and (f). CV, ISI, and firing rate distributions in (b,c,d) computed over 30 simulated networks, for experimental data over 489 trials. CV shows all data points (grey circles). CV median (white circle), interquartile range (black box), range (black line), kernel density estimate (violins) computed with data points *>* 3 std. devs. removed. Parameter scan results in (e,f) averaged over 5 simulated networks. Smoothing and contour lines added for visualization.

Both models were able to reproduce the activity statistics and the results were consistent over a range of refractory period distributions. Fig. 4e,f depict the difference between the models and experimental data when comparing the interspike interval distributions. The pattern of results demonstrate that heterogeneity is required to match the experimental data (purple region, white contours). When heterogeneity was low (*σ*_*ref*_ small), the models and experimental data were not in agreement. However, when refractory behavior was heterogeneous (*σ*_*ref*_ large enough), the models and experimental data matched well and this was true for a range of refractory period distributions (*µ*_*τ*_, *σ*_*τ*_). While both models fit the data, notably the anisotropic network did so for a larger set of parameters (larger purple region, see contours) and for larger means *µ*_*τ*_ and standard deviations *σ*_*τ*_ (purple region and contours shifted up and to the right, compare Fig. 4f vs. 4e) meaning wider distributions with more heterogeneity in refractory behavior across neurons (Fig. 4a).

Fig. 4b,c,d shows an example fit (gold stars, Fig. 4e,f) for the isotropic and anisotropic models for the coefficient of variation, ISI distribution, and firing rate distribution. Neurons recorded from M1 and PMd have more variability in their firing patterns than a Poisson process, evidenced by a coefficient of variation larger than one. Both the isotropic and anisotropic models matched this spiking behavior (Fig. 4b). Likewise, the models reproduced the ISIs (Fig. 4c) and firing rates (Fig. 4d) observed in the data and the fit was confirmed by comparing with a different M1/PMd recording (Supplementary Fig. S2).

### 2.3 Neuronal network activity resulting from symmetric and correlated asymmetric distance-dependent cortical connectivity

Next we were interested in the network activity of the isotropic and anisotropic models with parameters resulting from fitting both connectivity and activity statistics to cortical data. As discussed in the *Introduction*, while the isotropic network, with its symmetric connectivity, is expected to produce stationary activity bumps, the correlated asymmetric projections in the anisotropic network should give rise to spatiotemporal activity sequences. However, it remains unclear whether these findings hold true for the cortical parameter set.

Starting with the isotropic network model, we aimed to evoke intrinsically generated network activity under noisy background conditions. We conducted simulations in which a spatially localized burst of input was administered to the same network across 5 trials with different Gaussian white noise background input. The spatially localized burst varied on each trial, with a randomly selected 70% of the input region (404 of 576 neurons, a 24x24 neuron input region) receiving sufficient input current to spike at *t*_input_ = 10 ms. We repeated this for 10 different networks each generated based on the distributions for connectivity (*σ*_loc,exc_ = 0.14, *p*_con,rand_ = 0, see Fig. 2c,e and Fig. 3b,d) and activity (*µ*_*τ*_ = 7 ms, *σ*_*τ*_ = 5 ms, see Fig. 4e, gold star inside white contour) determined by the analyses in previous sections.

With symmetric distance-dependent connectivity and neuronal heterogeneity matched to experimental data, activity patterns in the isotropic network were dominated by spatially localized bumps (Fig. 5a,b). These activity bumps were transient with spatially local activity emerging and disappearing at different locations throughout the simulations (Fig. 5a,b). The pattern of transient emergence and disappearance was not consistent across trials as seen in the spike raster plots (Fig. 5a), activity plots of the two dimensional network topology (Fig. 5b), as well as when projected into low dimensional space (spanned by the first three principal components) with neural trajectories showing inconsistent behavior (Fig. 5c). Notably, the level of transience was variable across networks, with spatially localized activity at a particular location sometimes remaining relatively persistent throughout the 500 ms simulation (Supplementary Fig. S3).

**Figure 5.**
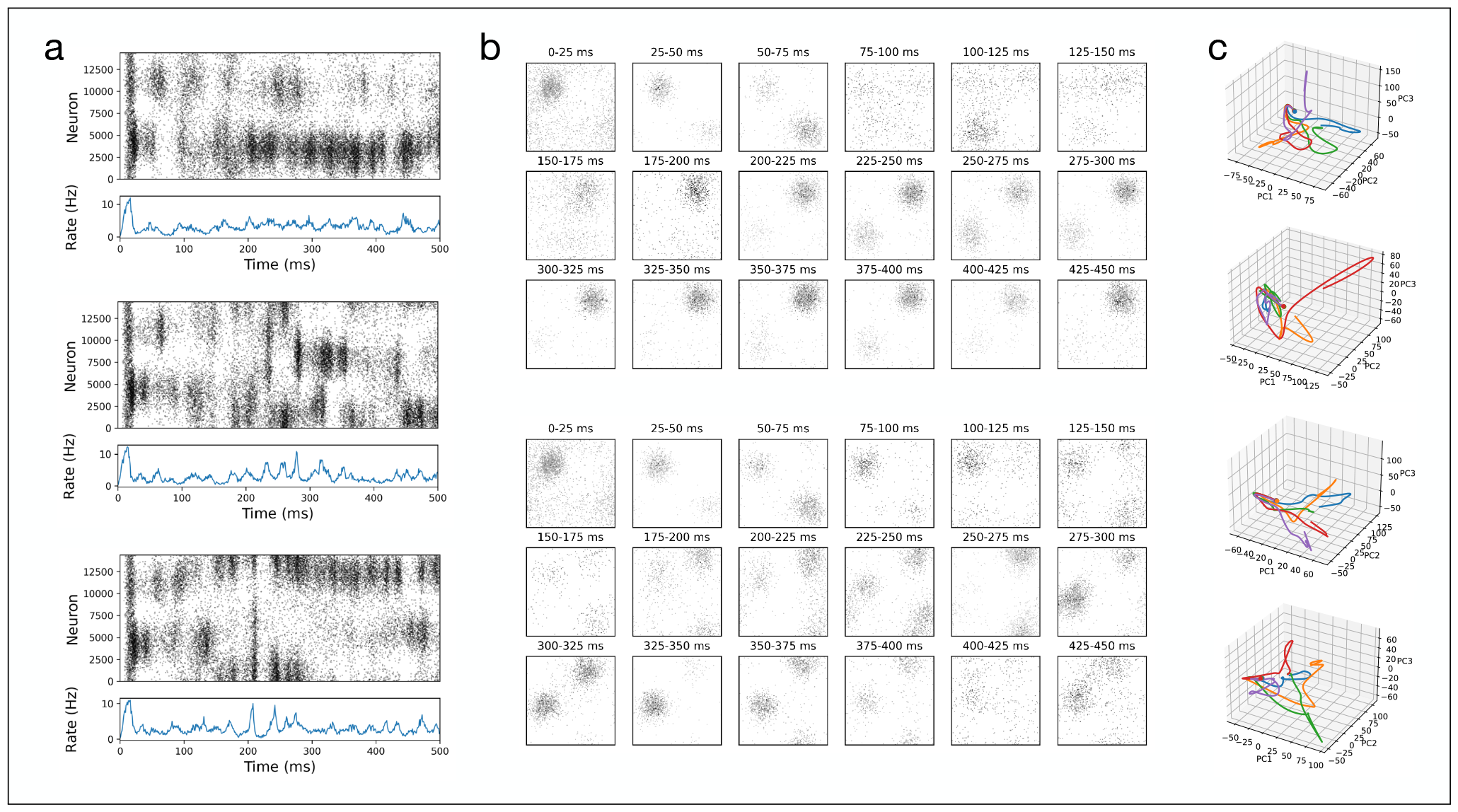
Transient spatially localized activity in networks with symmetric distance-dependent connectivity. **(a)** Three example trials depicting spiking behavior and average firing rates from the *N* = 14400 excitatory neurons in the isotropic network. **(b)** Activity over time organized in 2D according to neuron position shows transient bumps of activity appear and disappear. Top and bottom correspond to top and bottom in (a) respectively. Example 2 from (a) not shown. **(c)** Projection onto the first three principal components. Each 3D plot shows 5 trials from different network simulations. Data from (a) and (b) correspond to trials from the top panel in (c).

Next we turned to the activity statistics of the anisotropically connected network model, where we expect to find spatiotemporal activity sequences intrinsically generated by the correlated asymmetries in neuronal projections. To evoke intrinsically generated network activity we stimulated a spatially localized region (24x24 neurons) of the network across 5 trials with a randomly selected subset of the region receiving input (404 of 576 neurons, 70%) and different Gaussian white noise background input as before. We repeated this for 10 different anisotropically connected networks each generated based on the distributions for connectivity (*σ*_loc,exc_ = 0.14, *p*_con,rand_ = 0, see Fig. 2d,f and Fig. 3c,e) and activity (*µ*_*τ*_ = 6 ms, *σ*_*τ*_ = 15 ms, see Fig. 4f, gold star inside white contour) determined by the analyses in previous sections.

In contrast to the transient but stationary spatially localized activity in the case of symmetric distance-dependent connectivity, with correlated asymmetric projections we observed spatiotemporal sequences of activity (see Fig. 6). Activity patterns were again dominated by spatially localized bumps (Fig. 6a,b) which in this case moved throughout the network (Fig. 6a,b). In contrast with transient bumps in the isotropic case, spatiotemporal sequences showed consistency across trials, with visibly similar spike trains (Fig. 6a), similar movement of spatially local activity patterns (Fig. 6b), and accordingly, low dimensional neural trajectories that followed similar paths (Fig. 6c).

**Figure 6.**
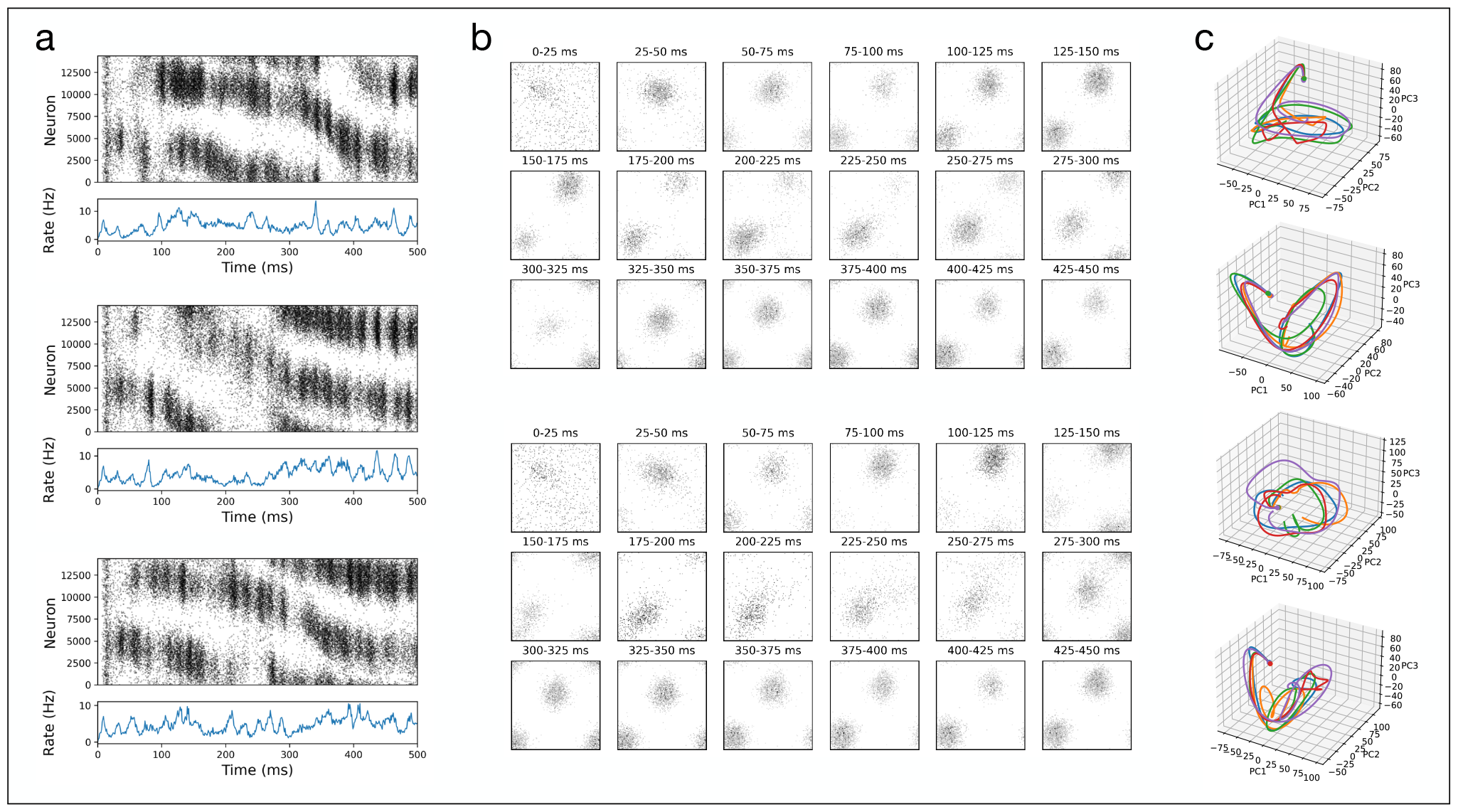
Spatiotemporal activity sequences in network with correlated spatial asymmetries in distance-dependent connectivity. **(a)** Three example trials depicting spiking behavior and average firing rates from the *N* = 14400 excitatory neurons in the anisotropic network. **(b)** Activity over time organized in 2D according to neuron position shows spatiotemporal sequences of activity moving through the network. Top and bottom correspond to top and bottom in (a) respectively. Example 2 from (a) not shown. **(c)** Projection onto the first three principal components. Each 3D plot shows 5 trials from different network simulations. Data from (a) and (b) correspond to trials from the top panel in (c).

Like the isotropic case, there was however variability across network simulations. We also observed transient spatiotemporal sequences in which the spatially localized input triggered a reliable initial response on the order of hundreds of milliseconds, after which point the activity across trials diverged (Fig. 7). Fig. 7a shows a network simulation in which neural trajectories in 3D principal component space are tightly aligned for the first 200 − 300 ms after stimulation, before beginning to diverge. The transient response and divergence can be clearly observed in behavior of the bumps of activity moving through the two dimensional network topology (Fig. 7c).

**Figure 7.**
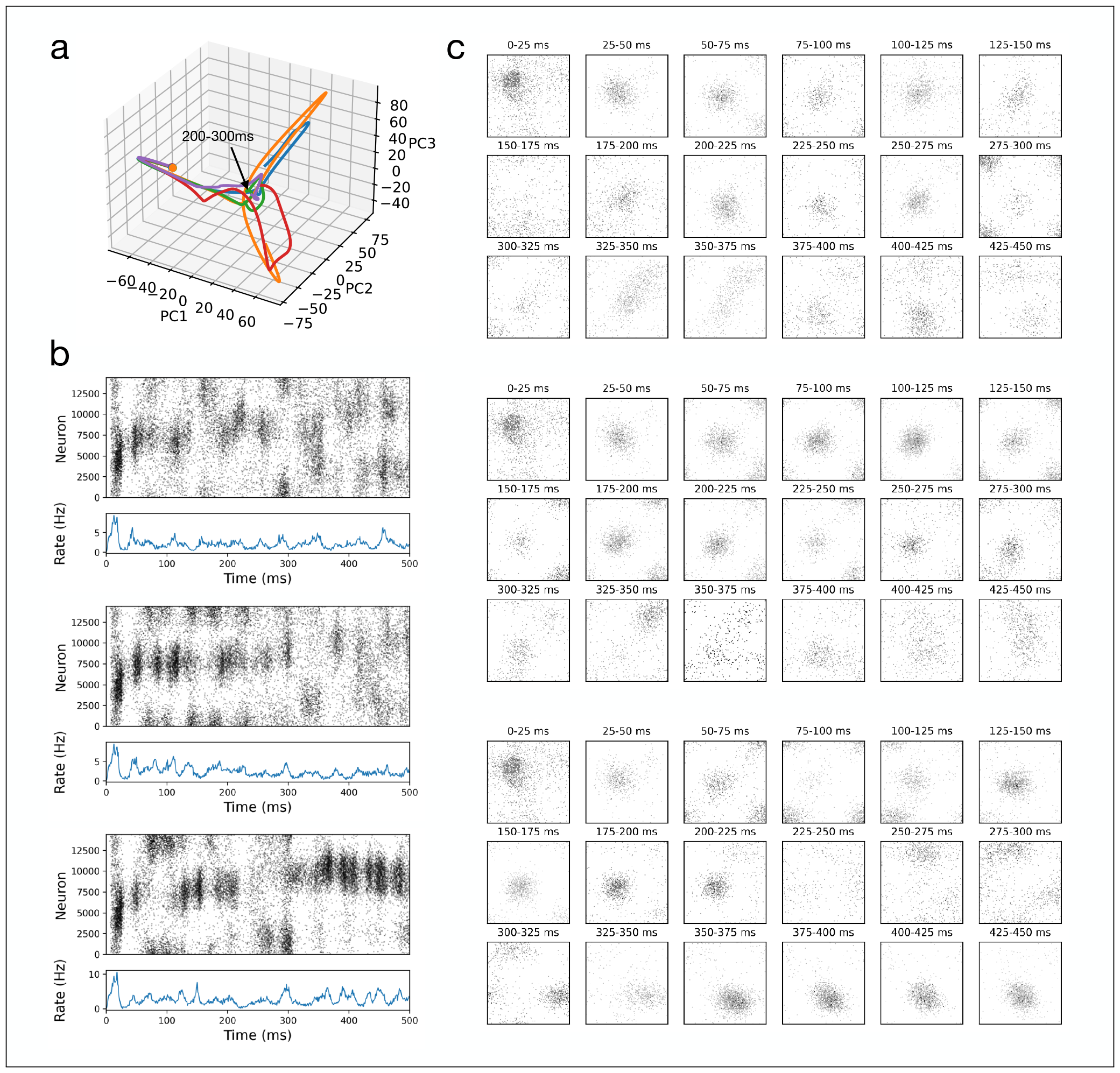
Transient spatiotemporal sequences arise with correlated asymmetric connectivity. **(a)** Projection of neural activity from five trials onto the first three principal components. Start of trial marked by dot. Trajectories follow the same path until they begin to diverge between 200 − 300 ms (black arrow). **(b)** Three example trials depicting spiking behavior and average firing rates from the *N* = 14400 excitatory neurons in the anisotropic network during a transient spatiotemporal sequence (corresponding to green, purple, and red lines, respectively). **(c)** Activity over time organized in 2D according to neuron position. Top and bottom correspond to top and bottom in (b) respectively. A spatiotemporal sequence of activity moves slowly through the network with activity patterns becoming visibly different between 225 − 300 ms.

## 3 Discussion

Here we have shown that recurrent network models with distance-dependent connectivity match both microcircuit connectivity and activity statistics of cortical data from rat and monkey. Symmetric distance-dependent connectivity was sufficient to explain the pattern of two- and three-neuron connectivity motifs observed in rat visual cortex^74^ (see also Miner and Triesch^80^). Surprisingly, introducing the assumption of asymmetric projections correlated in nearby neurons left the results unchanged, still fitting the nonrandom connectivity motifs. Likewise, single neuron activity statistics such as the distribution of firing rates, interspike intervals, and the coefficient of variation observed in monkey motor and premotor cortex could also be reproduced by both networks with distance-dependent symmetric as well as correlated asymmetric connectivity when heterogeneity in neuronal timescales was introduced via a distribution of refractory periods^79^.

With connectivity and neuronal timescales fitted to cortical data, both models produced spatially localized activity patterns in the 2D network topology. Symmetric connectivity gave rise to transient spatial patterns that emerged and disappeared, while correlated asymmetric connectivity produced spatiotemporal sequences of neuronal activity with spatially localized patterns becoming “unstuck in space”, and moving throughout the network.

Matching neuronal activity statistics from monkey motor/premotor cortex required different distributions of timescales depending on network connectivity. When connectivity was asymmetric and correlated, a wider distribution of timescales better fit the data (Fig. 4a). This result suggests that distributions of intrinsic timescales and refractory behavior may differ in brain regions that intrinsically generate spatiotemporal sequences compared with those that produce local transients. Notably, rhythmicity in firing rates emerged with heterogeneous refractory periods (Figs. 5-7), but how this relates to the precise shape of the refractory period distribution as well as the network level heterogeneity remains an open question.

Different to Spreizer et al.^37^, in the anisotropic network the parameters from fitting cortical data introduce a larger degree of variability in the network dynamics observed, with persisting spatial patterns, transient spatiotemporal sequences, and perturbations with spatiotemporal sequences briefly following different paths on different trials (Figs. 5, 6, 7 and Supplementary Figs. S3, S4, S5). Given the multiple sources of noise, with variability in the spatially localized input (404 of 576 neurons, or 70% of the input region randomly receiving input on a given trial) as well as background noise, differences in network behavior are perhaps not surprising. Whether the source of variability lies in particulars of the network topology remains to be explored. Equally interesting may be the effect of plasticity, in particular for task learning, on the different types of observed network behavior.

The focus of our work was motivated by experimental data pointing to spatial asymmetries in distance-dependent cortical connectivity^42–51^, and past theoretical work linking these connectivity structures to sequence generation^37^. However there are a number of different accounts of sequence generation in recurrent networks. Proposals involve continuous attractors^32,40^, discrete attractors or cell assemblies connected in feedforward chains^34,35,39^, as well as symmetric distance-dependent connectivity with adaptation mechanisms^28,30,32,36,81^ or distance-dependent transmission delays^70^ enabling localized activity to propagate.

Our work lends further support to the notion that locally connected recurrent network models offer a parsimonious account of transient attractor-like activity as well as spatiotemporal sequences while requiring only few experimentally supported assumptions. Correlated asymmetries in the projection patterns of nearby neurons may be a key feature of cortical networks, enabling intrinsic generation of spatiotemporal activity sequences based on locally defined flows to produce rich and stable dynamics.

## 4 Materials and Methods

### 4.1 Software

Simulations were conducted with a custom software framework written in Python. The anisotropic network model code was based on an implementation by Leo Hiselius^82^ in Brian2^83^ with extended simulation management functionality inspired by the architecture from *PeleNet*^84^.

### 4.2 Neuron and synapse model

The model is based on Spreizer et al.^37^ with a number of changes described below, see ref.^37^ for additional information. The subthreshold membrane potential *v*(*t*) of a neuron in the network is modeled as

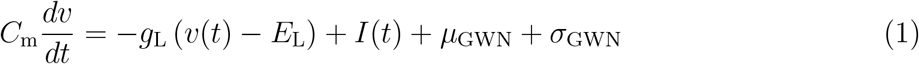

with the leak conductance *g*_L_, the membrane capacitance *C*_m_, the incoming current *I*(*t*), the leak potential *E*_L_, and Gaussian white noise with mean *µ*_GWN_ and standard deviation *σ*_GWN_.

Synapses between neurons are described by the transient current *I*_syn_, which is elicited by each presynaptic spike as

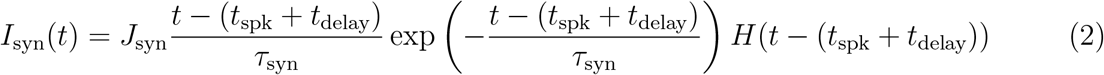

with *J*_syn_ the synaptic strength, *t*_spk_ denotes the time of a spike in the presynaptic neuron, *τ*_syn_ is the time constant of the synapse, *t*_delay_ is the delay between presynaptic spike and post-synaptic response, and *H* is the unit step function. The incoming synaptic current *I*_i_(*t*) of neuron *i* at time *t* is the sum over all the synaptic currents *I*_syn_(*t*) that the neuron receives from its *N*_syn_ synapses at time *t*:

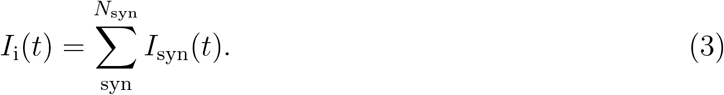

If *v*(*t*) reaches the threshold *V*_t_, a spike event occurs at time *t*_spk_. Spikes are transmitted from pre- to post-synaptic neuron with delay time *t*_delay_. After this event the neuron experiences an absolute refractory period, *τ*_ref_. Following the spike the neuron’s potential is reset to the resting potential *V*_r_. A spatially localized region of the network additionally receives a transient input burst described below (see *Methods: Spatially local input*).

### 4.3 Neuron subpopulations and spatial organisation

We model a network of excitatory and inhibitory neurons (see EI-network, Spreizer et al.^37^), with a ratio of four excitatory neurons to one inhibitory neuron based on cortical networks^85^.

When a presynaptic neuron is part of the excitatory subpopulation, *J*_syn_, defined in Equation 3, is equal to *J*_exc_ *>* 0, with excitatory synaptic strengths lognormally distributed (see *Methods: Lognormal synaptic weight distribution*). For inhibitory presynaptic neurons *J*_syn_ is equal to *J*_inh_, defined as:

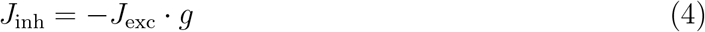

with *g* being the strength of the inhibition. Both subpopulations of neurons are homogeneously distributed on a square grid (x, y ∈ [0, 1]) with periodic boundaries (i.e. the surface of a toroid).

### 4.4 Interneuronal connectivity

Neurons form distance-dependent connections following a Gaussian distribution with a standard deviation *σ*_*loc,exc*_ for the projections of excitatory neurons and *σ*_*loc,inh*_ for the projections of inhibitory neurons. In the *anisotropic* network, for connections between excitatory neurons the center of this distribution is shifted away from the position of a neuron within the network, into a particular direction. This leads to a preferred axon direction *ϕ*_*i*_ for each neuron. The magnitude of the shift is determined by the shift *d* and is equivalent for all neurons. For the *isotropic* network, *d* = 0.

The direction of the shift *ϕ*_*i*_ is chosen individually for each neuron with Perlin noise, a gradient noise algorithm^37,86^. With this method, correlated directions are assigned to neurons that are close to each other. The result is an embedded local feedforward structure. Readers are referred to previous publications for further details^37,38,86^. Note that autapses are not permitted.

### 4.5 Lognormal synaptic weight distribution

In previous work, synaptic weights were equal for every synapse^37,38^. Here we assume a lognormal distribution of synaptic weights^74,87,88^. Weights are defined based on the distribution

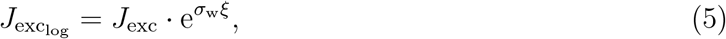

with *ξ* being a normally distributed variable and *J*_exc_ being the original constant weight described in *Methods Neuron subpopulations and spatial organisation. σ*_w_ is a prefactor that scales the influence of the random variable. By setting *µ* = ln(*J*_exc_), the resulting probability density function of the weights, *p*(*J*), is defined as (see Fukai et al.^89^, Michaelis^90^)

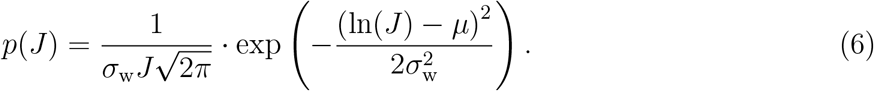

Note, that *µ* and *σ*_w_ are not the standard deviation and the mean of this distribution of *J* but rather of the distribution ln(*J*), since ln(*J*) is normally distributed.

### 4.6 Spatially local input

A rectangular subset of the neuronal grid containing *N*_g_ excitatory neurons received spatially local input^38^. In particular, only a predefined share *N*_s_ ≤ *N*_g_ of the neurons within this subset is stimulated with an instantaneous incoming current, strong enough to elicit a spike event at time *t*_input_. This ignores temporal dynamics in the transient input burst.

The spatially local input arrived at a region of *N*_*g*_ = 576 neurons (24x24 grid) of which on any given trial 70% (*N*_*s*_ = 404) received input current leading them to spike at time *t*_input_ = 0.01*s*. For simulations with multiple trials (Figs. 5-7 and Supplementary Figs. S3-S5), the subset of *N*_*s*_ = 404 neurons receiving input was randomly selected from the 24x24 neuron input region.

### 4.7 Distribution of refractory periods

Refractory periods were drawn from a Gaussian distribution

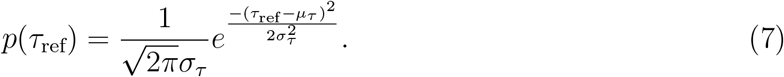

Here *σ*_*τ*_ is the standard deviation and *µ*_*τ*_ is the average of the distribution of *τ*_ref_. The probability that a neuron has a refractory period equal to *τ*_ref_ is, therefore, *p*(*τ*_ref_). Different locations of stimulation within a single neuron can lead to different refractory periods^79^, but for this work each neuron’s refractory period remains fixed.

We considered refractory period distributions with *µ*_*τ*_ ∈ [2 *ms*, 20 *ms*] and *σ*_*τ*_ ∈ [0 *ms*, 20 *ms*]. To exclude the possibility of negative and very small refractory periods the Gaussian distribution is cut at 1.5 milliseconds and capped at 120 milliseconds to avoid arbitrarily large *τ*_*ref*_ . We performed a parameter search over the mean and standard deviation of the refractory period distribution to fit single neuron activity statistics of the experimental data (Fig. 4 and Supplementary Fig. S2).

### 4.8 Connectivity motifs

We compared the recurrent connectivity in the *anisotropic* and *isotropic* network models to data from layer 5 pyramidal neurons in rat visual cortex^74^. Song et al.^74^ considered connectivity motifs of pairs and triads of recorded neurons and found the distributions differed from what is to be expected in a randomly connected network. For neuronal pairs there are three possible motifs, namely they can be connected bidirectionally, unidirectionally or not connected at all (Fig. 2a). For groups of three neurons, there are 16 different possible motifs (Fig. 3a). In the nomenclature of the three neuron motifs, the first digit represents the overall bidirectional connections, the second digit represents unidirectional connections and the third missing connections respectively. D, U and C refer to certain patterns arising from the directionality of the connections.

For the model, the probability *p*_con_ = 0.116 of a connection between two neurons is chosen such that it equals the experimental finding in Song et al.^74^. This probability is used in the synaptic initialisation processes of both excitatory and inhibitory neuron subpopulations.

Based on the connection probability, Song et al.^74^ calculated the probability of the different motifs under the assumption that the network is randomly connected. They compared these expected values to the experimentally observed motif counts and found a pattern of over- and underrepresentations of single motifs in the network beyond what is expected in a randomly connected network. Here we applied the same comparison to the motifs of synaptic connections in the model, which result from the connectivity generating rules described in *Methods: Interneuronal connectivity*. To determine the role of the anisotropy of the models synaptic connections, we evaluated the same statistics for a model without a shift in its connectivity (i.e. *d* = 0, see *Methods: Interneuronal connectivity*), the isotropic model. In this version the locality of the connectivity is preserved but neurons project symmetrically in all directions, thus lacking correlated anisotropy.

To quantify the distribution of motifs we split all neurons of the respective model randomly into pairs and counted the respective directed connections using their weight matrices. We repeated this procedure for neuronal triads. Every neuron was sampled only once for a pair and once for a triad.

We computed two and three neuron motif distributions as a function of two parameters. The connection locality *σ*_*loc,exc*_, i.e. the standard deviation of the Gaussian kernel for excitatory connections, was varied in the range *σ*_*loc,exc*_ ∈ [0.01, 0.25] and the proportion of random connections *p*_con,rand_ was varied in the range *p*_con,rand_ ∈ [0, 0.25]. Random connections are those not restricted to the local neighborhood of a neuron defined by *σ*_*loc,exc*_. The random connections replace local connections, hence the overall connection probability remains preserved. Note *p*_con,rand_ = 0 is a purely locally connected network and e.g. *p*_con,rand_ = 0.25 means that 25% of connections are randomly distributed and 75% of connections are local.

In the first step of the motif evaluation we estimated the occurrence *n*_occ_ of each motif (3 for two neuron motifs, 16 for three neuron motifs) within the respective number of neuron groups with size *n*_group_. With 14400 excitatory neurons in the model, sampling all neurons leads to *n*_group_ equalling 7200 samples for neuron pairs and 4800 samples for triads of neurons. We then calculated the relative representation *r*_rel_ for each motif by dividing the observed occurrence *n*_occ_ by its expected occurrence under the assumption of a random connectivity, as it was performed by Song et al.^74^. The expected occurrence can be determined by the overall number of analysed neurons groups *n*_group_ times the probability *p*_motif_ of the motif in a randomly connected network

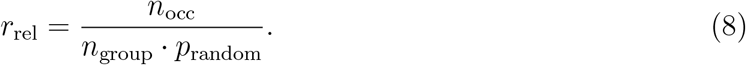

The probabilities *p*_motif_ for the different motifs are listed in Table 1 for two-neuron motifs and Table 2 for three-neuron motifs. They depend on the overall connection probability of two neurons *p*_con_ equal to 11.6% in the model, which is the connection probability from the experimental data^74^.

**Table 1.**
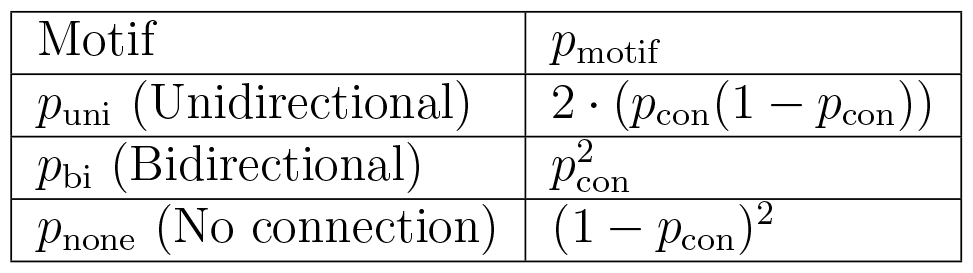
Probabilities of two neuron motifs expected in a randomly connected network.

| Motif | $p_{\text{motif}}$ |
| --- | --- |
| $p_{\text{uni}}$ (Unidirectional) | $2 \cdot (p_{\text{con}}(1 - p_{\text{con}}))$ |
| $p_{\text{bi}}$ (Bidirectional) | $p_{\text{con}}^2$ |
| $p_{\text{none}}$ (No connection) | $(1 - p_{\text{con}})^2$ |

**Table 2.** Probabilities of three-neuron motifs expected in a randomly connected network depending of the probabilities of two-neuron motifs *p*_uni_, *p*_bi_, *p*_none_.

| Motif | $p_{\text{motif}}$ |
| --- | --- |
| **012** | $6 \cdot (p_{\text{none}}^2) \cdot p_{\text{uni}}$ |
| **102** | $3 \cdot (p_{\text{none}}^2) p_{\text{bi}}$ |
| **021D** | $3 \cdot p_{\text{none}} \cdot p_{\text{uni}}^2$ |
| **021U** | $3 \cdot p_{\text{none}} \cdot p_{\text{uni}}^2$ |
| **021C** | $6 \cdot p_{\text{none}} \cdot p_{\text{uni}}^2$ |
| **111D** | $6 \cdot p_{\text{none}} \cdot p_{\text{uni}} \cdot p_{\text{bi}}$ |
| **111U** | $6 \cdot p_{\text{none}} \cdot p_{\text{uni}} \cdot p_{\text{bi}}$ |
| **201** | $3 \cdot p_{\text{none}} \cdot p_{\text{bi}}^2$ |
| **030T** | $6 \cdot p_{\text{uni}}^3$ |
| **030C** | $2 \cdot p_{\text{uni}}^3$ |
| **120D** | $3 \cdot (p_{\text{uni}}^2) \cdot p_{\text{bi}}$ |
| **120C** | $6 \cdot (p_{\text{uni}}^2) \cdot p_{\text{bi}}$ |
| **120U** | $3 \cdot (p_{\text{uni}}^2) \cdot p_{\text{bi}}$ |
| **210** | $6 \cdot p_{\text{uni}} \cdot p_{\text{bi}}^2$ |
| **300** | $1 \cdot p_{\text{bi}}^3$ |

**Table 3.**
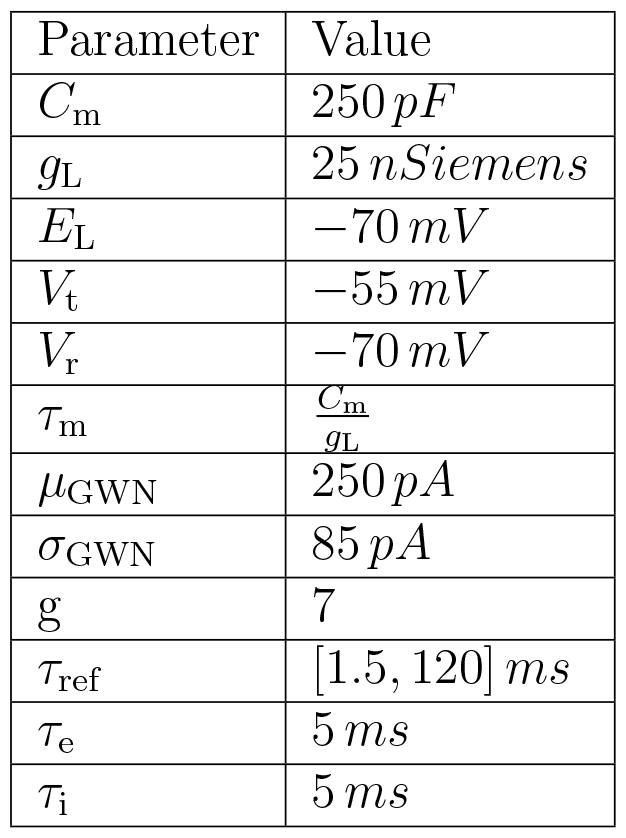
Neuron parameters.

**Table 4.** Synaptic parameters.

| Parameter | Value |
| --- | --- |
| $J_{\text{exc}}$ | $10 \cdot e\text{ pA}$ |
| $J_{\text{inh}}$ | $-g \cdot J_{\text{exc}}$ |
| $\sigma_{\text{loc,exc}}$ | $0.14$ |
| $\sigma_{\text{loc,inh}}$ | $\frac{\sigma_{\text{loc,exc}}}{0.75}$ |
| $t_{\text{delay}}$ | $1\text{ ms}$ |
| $p_{\text{conexc}}$ | $0.116$ |
| $p_{\text{coninh}}$ | $0.116$ |
| $s_{\text{perlin}}$ | $2$ |
| $d_{\text{shift}}$ | $1$ |
| $p_{\text{con,rand}}$ | $0$ |

**Table 5.** Network and simulation parameters.

| Parameter | Value |
| --- | --- |
| $t_{\text{simulation}}$ | $0.5\text{ s}$ |
| $t_{\text{input}}$ | $0.01\text{ s}$ |
| $n_e$ | $14400$ |
| $n_i$ | $3600$ |

To estimate *p*_motif_ for the three neuron motifs, Song et al.^74^ did not use *p*_uni_, *p*_bi_ and *p*_none_ as they would be expected in a random network, but rather the probability of occurrence of two-neuron motifs they observed for the rat visual cortex neurons. The reason is to ensure that overrepresented three-neuron patterns are not spuriously detected simply because they contain an overrepresented two-neuron pattern. We apply the same approach using *p*_uni_, *p*_bi_ and *p*_none_ from the experimental data^74^ .

For each parameter set (*σ*_*loc,exc*_, *p*_con,rand_), we estimated the average over *N*_seed_=5 different network initialisations generated by using different random seeds for the connectivity. In the anisotropic model every initialisation has a different landscape of preferred directions (see *Methods: Interneuronal connectivity*) as well as a different initial pseudorandom number generator state for the process of the determination of synaptic connections based on *p*_con_ within the model. Since there is no preferred direction in the isotropic model only the latter changes within its initialisations.

The mean relative representation 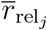 of a motif *j* for a certain parameter set is computed as

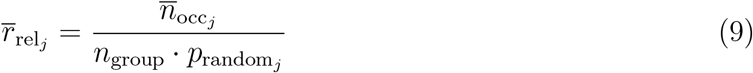

with 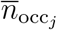, the mean occurrence of every motif, defined as

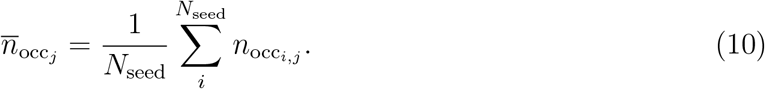

To estimate the deviation 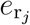 of the relative representation 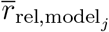 of a motif *j* from the model to the relative representation 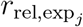 of the same motif *j* in the experimental data^74^, we calculated the absolute value of their difference and normalised it by the relative representation within the experimental data 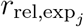

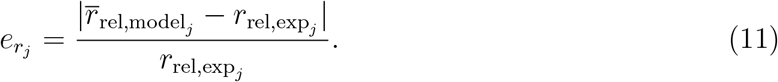

In order to have a measure to compare the match between the model motifs and the motifs from the experimental data for many sets of parameters, we averaged the 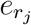 for the different motifs of two neuron motifs (*N*_motif_ = 3) as well as for three neuron motifs (*N*_motif_ = 16) giving the *mean relative error E*_*r*_, defined as:

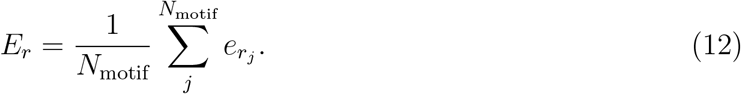

### 4.9 Electrophysiological data

Methodological details relating to recording, waveform identification, and offline sorting are provided in the methods of previous work^91^ for neural recordings from macaque monkey cortical areas PMd and M1 via floating microelectrode arrays (128-channels total; 64-channels per area). In this paper, we make use of similar data from a memory-guided center-out reach task to one of four directions (0°, 90°, 180°, 270°, pseudo-randomly interleaved from trial to trial), which was collected for another project (Nowak, Morel & Gail, unpublished).

Data from two different recording sessions (referred to as DS1 and DS2) were analyzed for reliability, the first with 66 neurons and the second with 56 neurons (in M1 and PMd combined). We considered only trials in which the monkey successfully performed the task for analysis (489 and 438 successful trials, respectively). For each trial, we considered spiking data in the time window from instruction to initiate the movement until the movement end(duration, mean *±* standard deviation for DS1: 575.60 *±* 76.05 milliseconds; DS2 568.18 *±* 89.65 milliseconds).

### 4.10 Neuronal activity measures

We compared single neuron activity statistics from the electrophysiological data to simulations of *aniosotropically* and *isotropically* connected network models in response to a transient spatially local burst of input (see *Methods: Spatially local input*) with background noise. Here we varied the mean and standard deviation of the refractory period distribution (see *Methods: Distribution of refractory periods*) and quantified firing rates, interspike intervals, and the coefficient of variation.

#### 4.10.1 Neuronal firing rates

The firing rate *ν*_i_ of a neuron *i* is computed by dividing the number of spikes *n*_spk_ by the evaluated time interval *T*

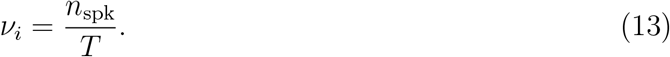

We compared the distribution of firing rates in simulations with the experimental data (see *Methods: Comparison of distributions*).

#### 4.10.2 Interspike intervals

An interspike interval (ISI) measures the time interval *s*_*k*_ in between two subsequent spikes spk_*k*_ and spk_*k*+1_ of a neuron:

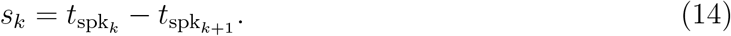

We quantified the distribution of ISIs across neurons. Details on this analysis can be found in *Methods: Comparison of distributions*.

#### 4.10.3 Coefficient of variation

The coefficient of variation (*C*_V_) measures the regularity of a spike train compared to a spike train that would arise from a Poisson process^92^. In a Poisson process neurons have a constant firing rate *ν* and spikes occur independently and stochastically^92^, based on *ν*. Therefore the average number of spikes in the interval T is *ν* · *T* . The coefficient of variation for a Poisson process is equal to 1 (*C*_V_ = 1). It holds that *C*_V_ *<* 1 if the spike train is more regular and *C*_V_ *>* 1 if it is less regular than a Poisson process^92^. *C*_V_ is estimated by the division of the standard deviation of ISIs, *σ*_ISI_, of a spike train by the expected value of the ISIs, *µ*_ISI_, of the same spike train

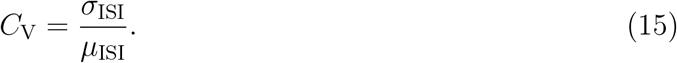

### 4.11 Comparison of distributions

#### 4.11.1 Binning of ISIs and firing rates

We binned and normalized the ISIs and firing rates for comparison between experimental and simulated data. For ISI distributions, data was binned into *N*_bin_ = 100 bins covering a range of ISIs between 0 ms and 200 ms and the width of each bin was *w*_b_ = 2 milliseconds. For the firing rates, *N*_bin_ = 50 bins were calculated ranging from 0 Hz to 100 Hz with *w*_b_ = 2 Hz. Neurons with less than one spike in a trial were not considered for firing rate distributions and those with less than 2 spikes were not considered for ISI distributions.

For the experimental data, in DS1 *N*_sample_ = 66 neurons were recorded over *N*_trials_ = 489 trials and in DS2 *N*_sample_ = 56 neurons were recorded over *N*_trials_ = 438 trials. Bins are normalised by dividing them by the number of neurons times the number of trials.

In the simulation data, for each simulation we sampled *N*_sample_ = 2000 neurons. Fig. 4b-d and Supplementary Fig. S2a-c show binned data for *N*_seed_ = 30 simulations and the comparisons in Fig. 4e,f and Supplementary Fig. S2d,e are averages over *N*_seed_ = 5 simulations. This means for the former, ISIs and firing rates from the *N*_seed_ = 30 were binned together and normalized by *N*_sample_ · *N*_seed_, and for the latter each simulation was normalized separately and then averaged.

#### 4.11.2 Differences between distributions

To estimate the differences between model and experimental distributions we calculated the differences *d*_*i*_ of every bin 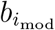 in the model to its counterpart in the experimental data 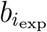. The absolute value of the difference is taken to compare the magnitude of the deviation. The difference measure is given by

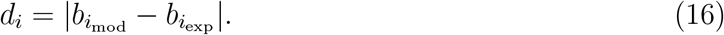

The *N*_bin_ bin-differences of a single network initialisation are summed up and an average of this sum is taken from the five initialisations. This results in the mean distribution difference of ISI distributions 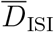

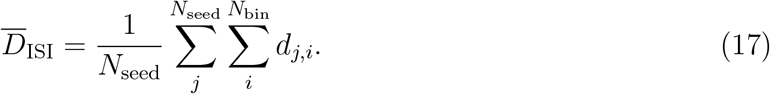

The comparison of ISI distributions between the models and DS1 is shown in Fig. 4e,f and with DS2 in Supplementary Fig. S2d,e.

### 4.12 Stimulus-evoked activity

For the stimulus-evoked activity, neurons were initialized at their rest potential and an instantaneous spatially localized input was administered at *t* = 10 ms during ongoing background input (see *Methods: Spatially local input*). Simulations lasted 500 ms with time step 0.1 ms. For activity statistics comparisons, we repeated the stimulus-evoked measurements for five different network initialisations for Fig. 4e,f and Supplementary Fig. S2d,e and for 30 initialisations for Fig. 4b-d and Supplementary Fig. S2a-c.

To assess transient bump vs. spatiotemporal sequences, we considered 10 different networks initialisations. In this case, in the same network five trials were conducted with stimulation delivered to a randomly selected 70% of the input region (404 of 576 neurons, a 24x24 neuron input region) as described before.

#### 4.12.1 Spike rasters

Spike raster plots show all spikes for all excitatory neurons with neuron order simply being the 2D network topology unrolled into a 1D vector.

#### 4.12.2 Instantaneous firing rates

Instantaneous firing rate plots were made by temporally binning spikes from all *n*_*e*_ = 14400 excitatory neurons into 1 ms bins and dividing the number of spikes in a temporal bin *i, n*_*spks*_(Δ*t*_*i*_), by the bin size Δ*t*_*i*_ and number of neurons *n*_*e*_. In a formula:

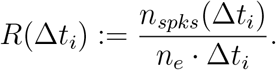

#### 4.12.3 2D activity plots

For 2D activity plots we simply plotted a black dot at the location in the 2D network topology for each spike that occurred within a given 25 ms temporal window.

#### 4.12.4 Principal component analysis

Spike data was binned into 1 ms bins resulting in a 14400x500 matrix containing spikes for *n*_*e*_ excitatory neurons for each time bin. The data was smoothed by convolving the spike train of each neuron with a Gaussian with temporal standard deviation 25 ms (using Python method `numpy.convolve` with `mode=“same”`). For the 5 trials from a single network, the smoothed spike rasters were concatenated into a 14400x2500 dimensional matrix. Each neuron’s smoothed spike train was centered and scaled to unit variance (using `sklearn.preprocessing.StandardScaler`). Principal component analysis was applied to the resulting matrix (using `sklearn.decomposition` PCA) and the smoothed, centered, and scaled spike raster was projected onto the first three principal components giving a matrix of size 3x2500. We separated the matrix back into the constituent trials (5 trials of 3x500) and plotted them in 3D.

### 4.13 Smoothing data for visualisation

For parameter search figures comparing model and experimental data for both the connectivity and activity statistics, we used a Gaussian filter with *σ* = 0.7 to smooth the data using `scipy.ndimagegaussian filter` and upsampled by a factor of 10 using `scipy.ndimage.zoom`.

### 4.14 Parameters

If not stated differently the following parameters of the model used in this work was defined as following. For the isotropic network model *d*_shift_ was set to 0.

## 5 Author contributions

Conceptualization ABL FE CM AK CT Data Curation ABL FE CM JN AG Formal Analysis ABL FE CM Funding Acquisition ABL AK AG CT Investigation ABL FE CM JN Methodology ABL FE CM Project Administration ABL CM Resources AG, CT Software FE ABL CM Supervision AK CT Validation ABL FE CM Visualization ABL FE CM Original Draft Preparation ABL FE Review & Editing ABL FE CM JN AG AK CT.

## 6 Acknowledgements

We thank Leo Hiselius for making his code publicly available and for discussions related to his code. We thank Benjamin Schulz for programming work related to the motif analysis. This work was supported by the German Research Foundation (Deutsche Forschungsgemeinschaft) in the context of the SFB 889 ‘Cellular Mechanisms of Sensory Processing’ (C4) and the European Commission in the context of the Plan4Act consortium (Grant H2020-FETPROACT-16 732266; project WP1), as well as the Swedish Research Council and StratNeuro. ABL is supported by a Natural Sciences and Engineering Research Council of Canada PGSD-3 scholarship.

## Supplementary Figures

**Figure S1.**
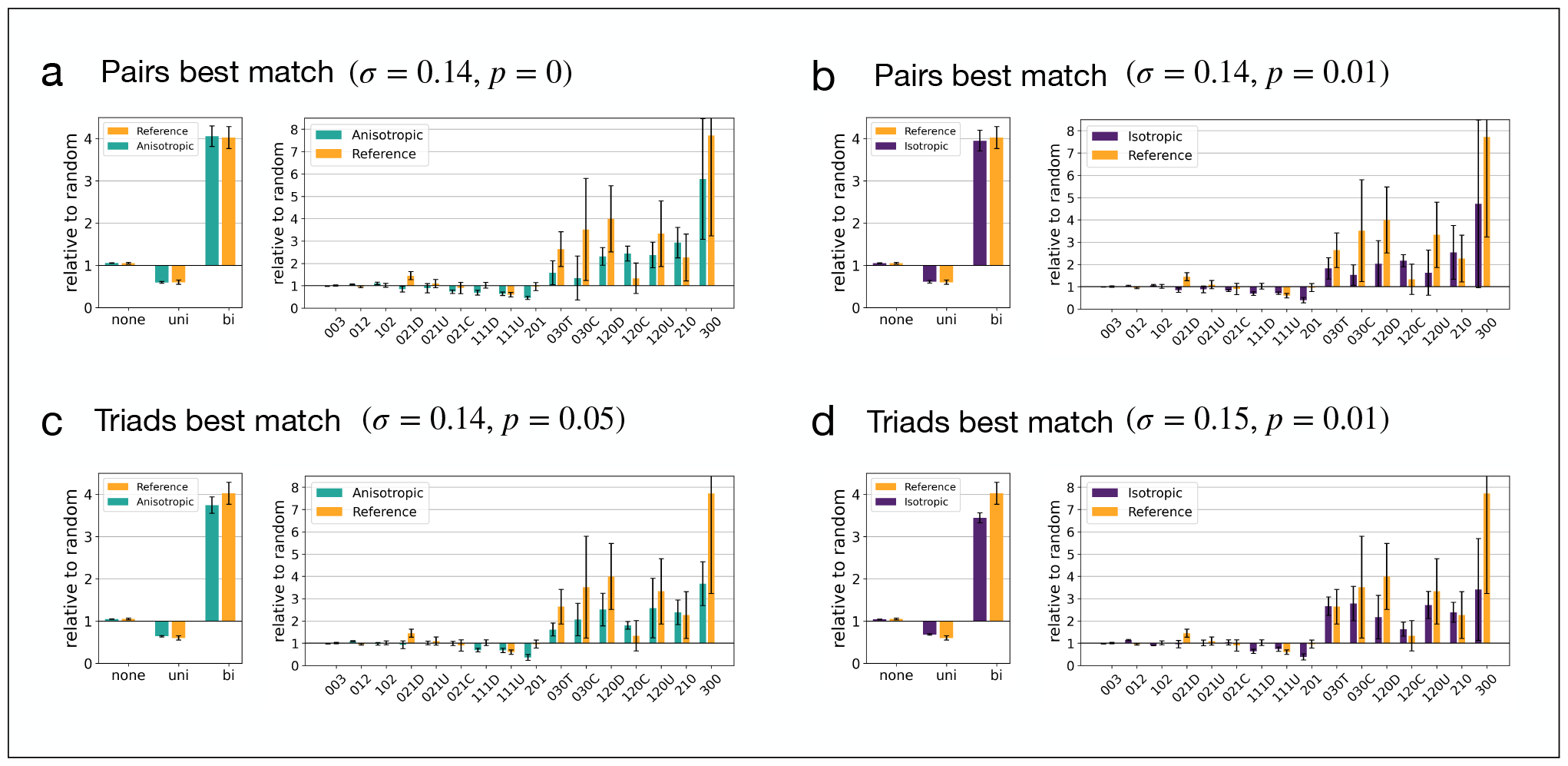
Two and three neuron motif counts for parameters best matching experimental data^74^ (related to Figs. 2 and 3). **(a)** Two and three neuron motif distributions for the anisotropic network model for the parameters that minimized the mean relative error between the isotropic network vs. rat visual cortex data for motif counts in neuron pairs (see Fig. 2d). **(b)** Same as (a) but for the isotropic network (see Fig. 2c). **(c)** Two and three neuron motif distributions for the anisotropic network model for the parameters that minimized the mean relative error between the isotropic network vs. rat visual cortex data for motif counts in groups of three neurons (see Fig. 2c). **(d)** Same as (c) but for the isotropic network (see Fig. 3b).

**Figure S2.**
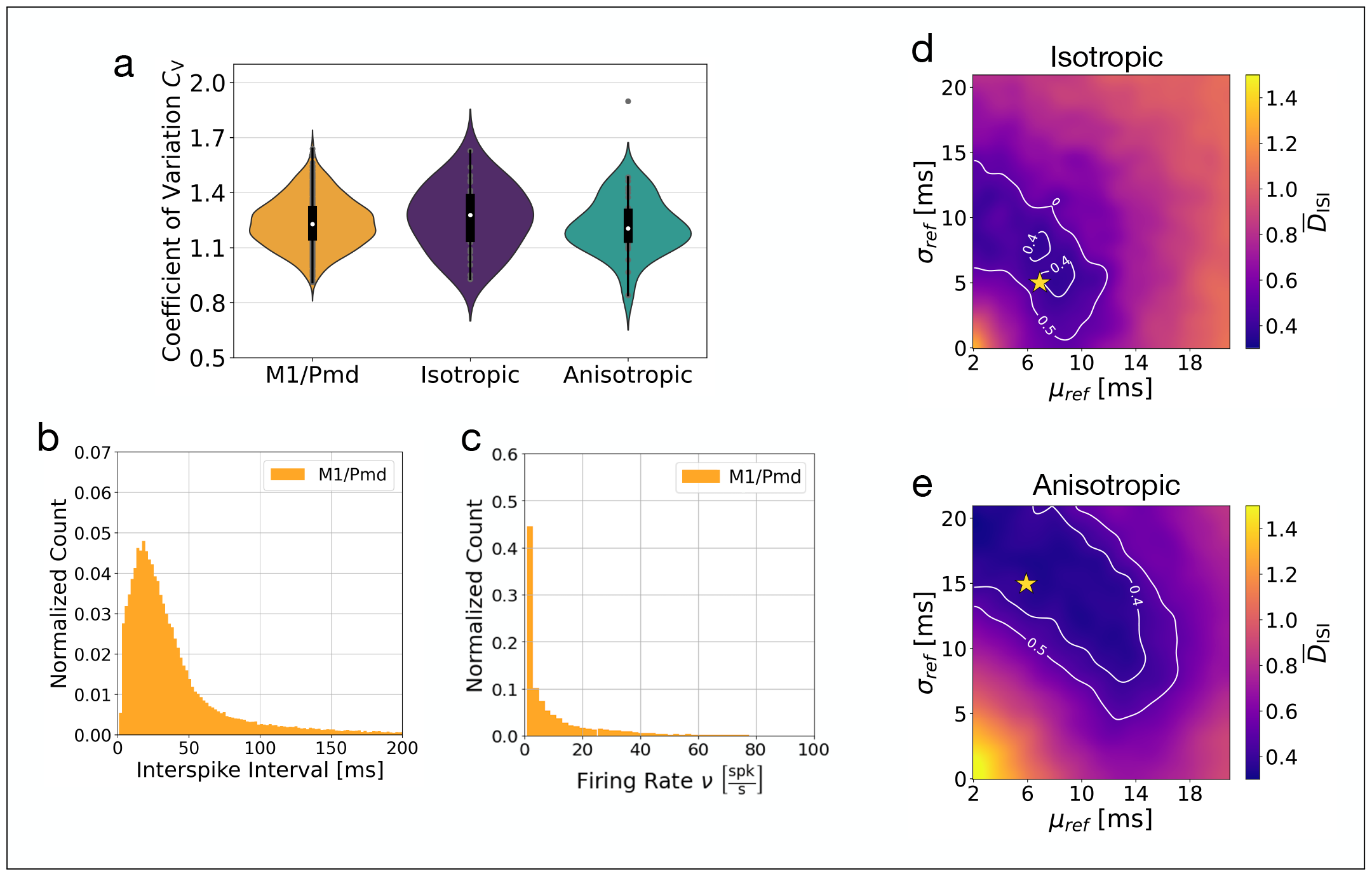
Single neuron activity statistics match to second M1/PMd dataset (related to Fig. 4). **(a)** Coefficient of variation for M1/PMd and the isotropic and anisotropic networks (reproduced from Fig. 4b). **(b)** ISI and firing rate distributions from the second monkey dataset. **(d**,**e)** Fits between data and models for isotropic and ansisotropic connectivity.

**Figure S3.**
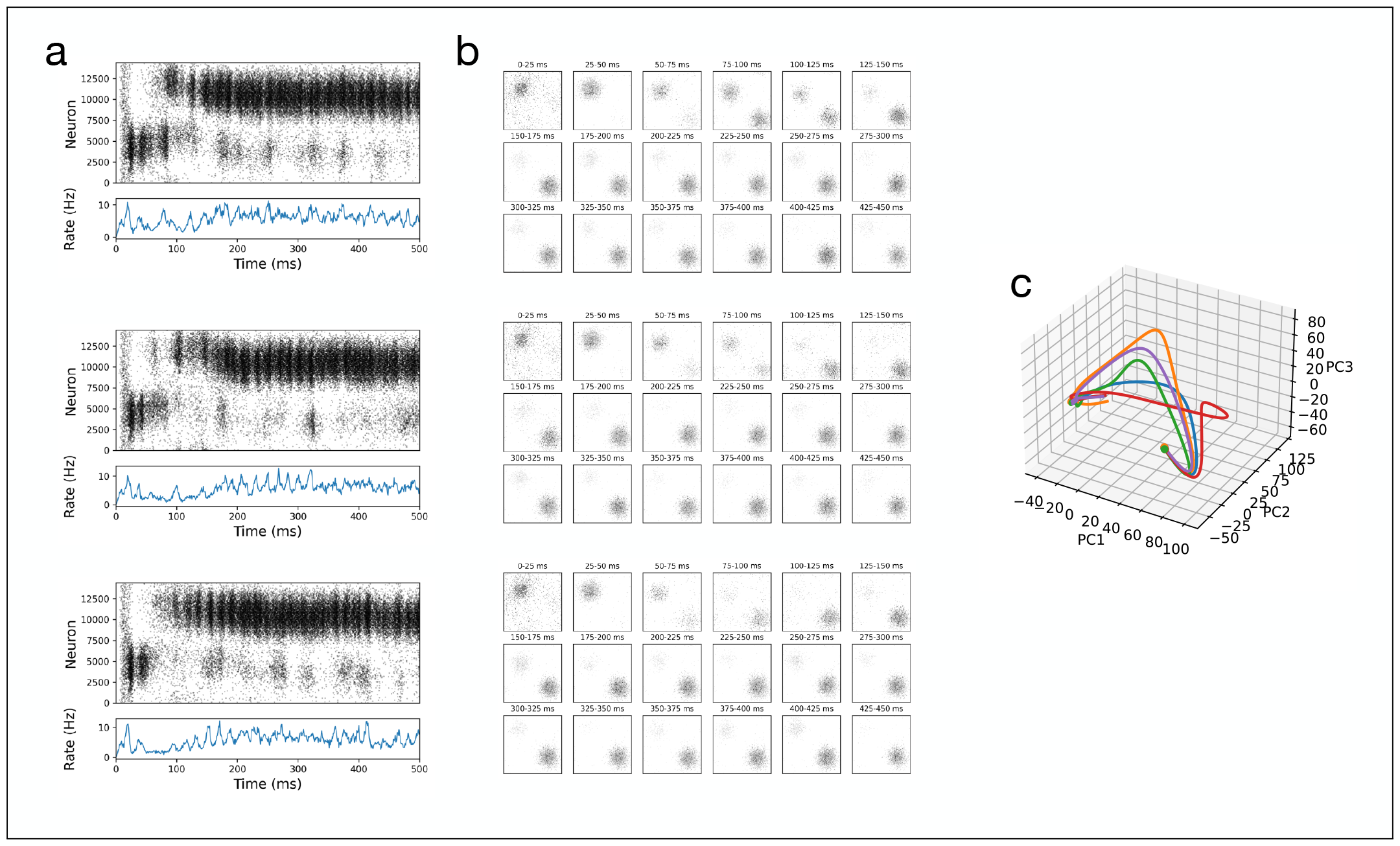
Persistent spatially localized activity in network with symmetric distance-dependent connectivity fit to experimental data (related to Fig. 5). **(a)** Spike rasters for three trials of one isotropic network simulation. **(b)** Activity over time organized in 2D according to neuron position shows bumps of activity. Bump persists at one location for a few hundred milliseconds. **(c)** Neural activity projected onto the first three principal components. Trajectories converge to one point in PC space and remain there, in agreement with the persistent bump of activity in the network.

**Figure S4.**
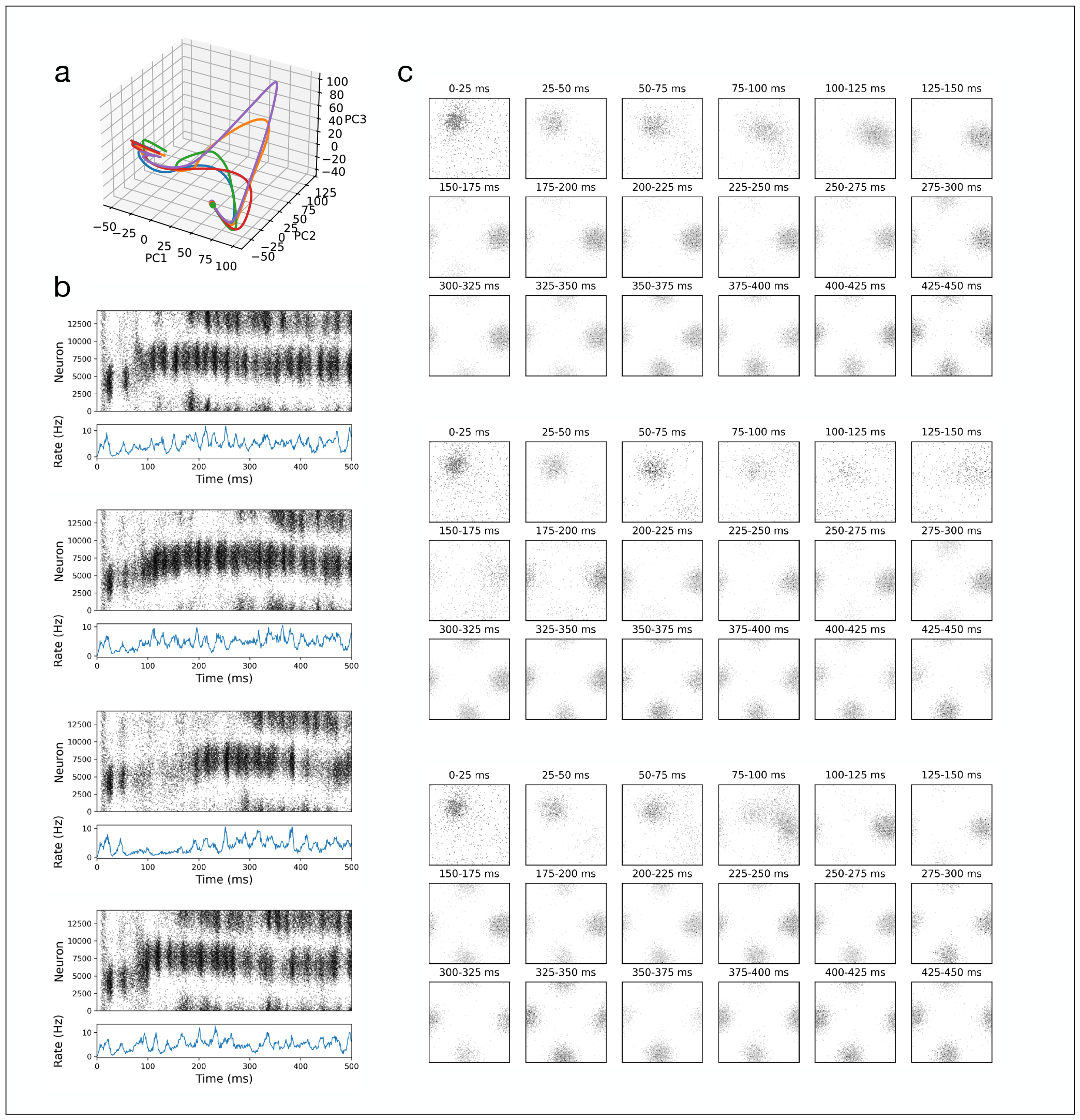
Trajectories taking different paths in anisotropic network (related to Fig. 6). **(a)** Neural activity projected onto the first three principal components. Trajectories for different trials take different paths related to the differences in spatiotemporal sequences **(b)** Spike rasters for four trials shows differences in activity in the early part of the trial before becoming similiar thereafter. **(c)** Activity over time organized in 2D according to neuron position shows bumps of activity which are visibly different in the early part of the trial, becoming similar as the trial progresses.

**Figure S5.**
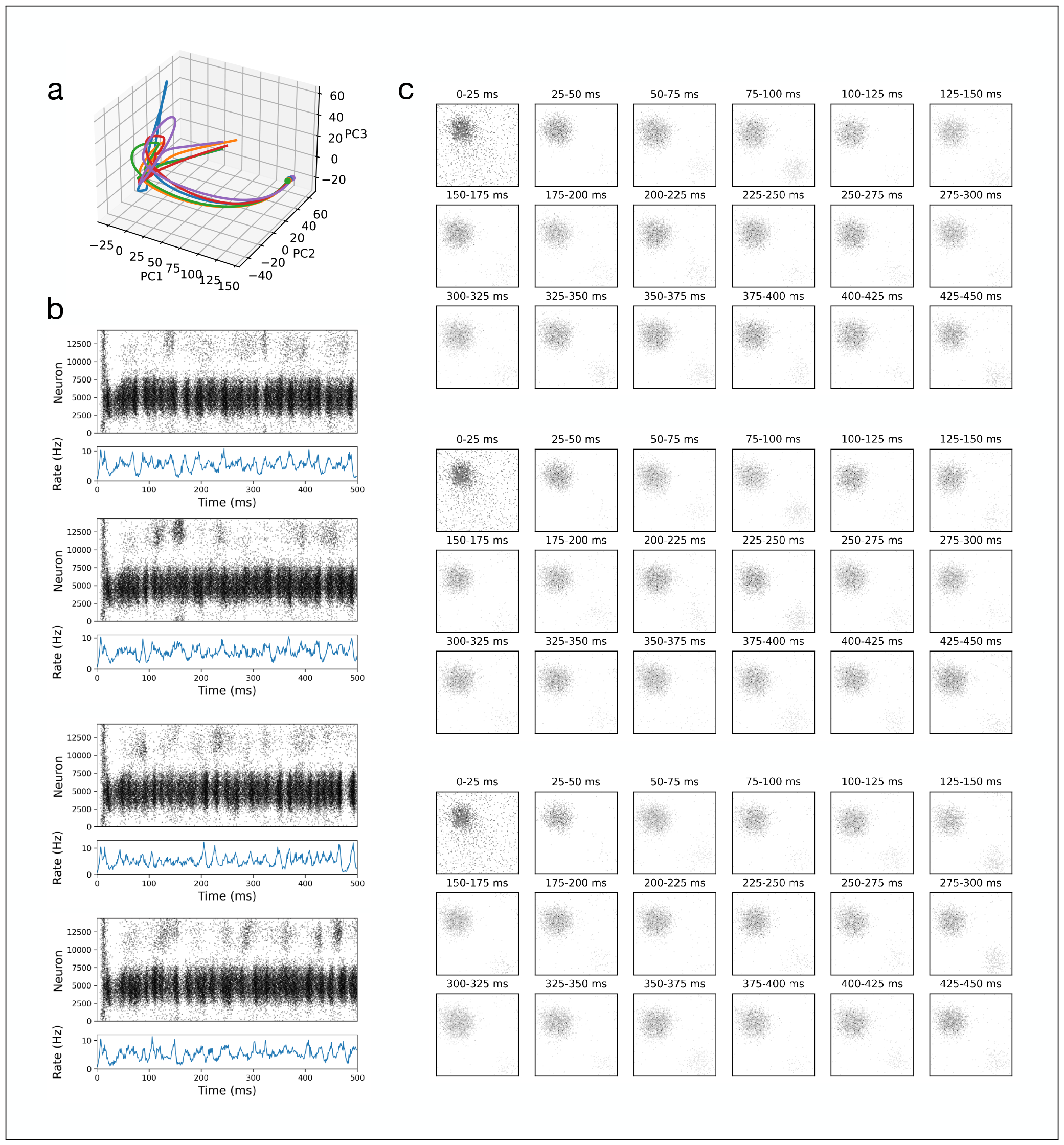
Persistent spatially localized activity in anisotropic network with correlated asymmetric distance-dependent connectivity fit to experimental data (related to Fig. 6). **(a)** Neural activity projected onto the first three principal components. Trajectory paths remain closely aligned and converge to one region in PC space in agreement with the persistent bump of activity in the network. **(b)** Spike rasters for four trials. **(b)** Activity over time organized in 2D according to neuron position shows bump persisting.

## References

[1] Christopher D Harvey, Philip Coen, and David W Tank. Choice-specific sequences in parietal cortex during a virtual-navigation decision task. Nature, 484(7392):62–68, 2012.

[2] Dezhe Z Jin, Naotaka Fujii, and Ann M Graybiel. Neural representation of time in cortico-basal ganglia circuits. Proceedings of the National Academy of Sciences, 106(45):19156–19161, 2009.

[3] Konstantin I Bakhurin, Vishwa Goudar, Justin L Shobe, Leslie D Claar, Dean V Buonomano, and Sotiris C Masmanidis. Differential encoding of time by prefrontal and striatal network dynamics. Journal of Neuroscience, 37(4):854–870, 2017.

[4] Christopher J MacDonald, Kyle Q Lepage, Uri T Eden, and Howard Eichenbaum. Hippocampal “time cells” bridge the gap in memory for discontiguous events. Neuron, 71(4): 737–749, 2011.

[5] Rainer W Friedrich and Gilles Laurent. Dynamic optimization of odor representations by slow temporal patterning of mitral cell activity. Science, 291(5505):889–894, 2001.

[6] Richard HR Hahnloser, Alexay A Kozhevnikov, and Michale S Fee. An ultra-sparse code underliesthe generation of neural sequences in a songbird. Nature, 419(6902):65–70, 2002.

[7] Henrik Lindén, Peter C Petersen, Mikkel Vestergaard, and Rune W Berg. Movement is governed by rotational neural dynamics in spinal motor networks. Nature, 610(7932):526–531, 2022.

[8] Juan A Gallego, Matthew G Perich, Stephanie N Naufel, Christian Ethier, Sara A Solla, and Lee E Miller. Cortical population activity within a preserved neural manifold underlies multiple motor behaviors. Nature communications, 9(1):4233, 2018.

[9] Jean-Baptiste Eichenlaub, Beata Jarosiewicz, Jad Saab, Brian Franco, Jessica Kelemen, Eric Halgren, Leigh R Hochberg, and Sydney S Cash. Replay of learned neural firing sequences during rest in human motor cortex. Cell Reports, 31(5):107581, 2020.

[10] William E Skaggs and Bruce L McNaughton. Theta phase precession in hippocampal. Hippocampus, 6:149–172, 1996.

[11] Eva Pastalkova, Vladimir Itskov, Asohan Amarasingham, and Gyorgy Buzsaki. Internally generated cell assembly sequences in the rat hippocampus. Science, 321(5894):1322–1327, 2008.

[12] György Buzsáki and David Tingley. Space and time: the hippocampus as a sequence generator. Trends in cognitive sciences, 22(10):853–869, 2018.

[13] John O’Keefe. Place units in the hippocampus of the freely moving rat. Experimental Neurology, 51(1):78–109, 1976.

[14] George Dragoi and György Buzsáki. Temporal encoding of place sequences by hippocampal cell assemblies. Neuron, 50(1):145–157, 2006.

[15] David J Foster and Matthew A Wilson. Reverse replay of behavioural sequences in hippocampal place cells during the awake state. Nature, 440(7084):680, 2006.

[16] Kamran Diba and György Buzsáki. Forward and reverse hippocampal place-cell sequences during ripples. Nature Neuroscience, 10(10):1241, 2007.

[17] Mehrab N Modi, Ashesh K Dhawale, and Upinder S Bhalla. Ca1 cell activity sequences emerge after reorganization of network correlation structure during associative learning. Elife, 3:e01982, 2014.

[18] Sen Cheng. The crisp theory of hippocampal function in episodic memory. Frontiers in neural circuits, 7:88, 2013.

[19] Upinder S Bhalla. Dendrites, deep learning, and sequences in the hippocampus. Hippocampus, 29(3):239–251, 2019.

[20] Soledad Gonzalo Cogno, Horst A Obenhaus, Ane Lautrup, R Irene Jacobsen, Claudia Clopath, Sebastian O Andersson, Flavio Donato, May-Britt Moser, and Edvard I Moser. Minute-scale oscillatory sequences in medial entorhinal cortex. Nature, pages 1–7, 2023.

[21] Artur Luczak, Peter Barthó, Stephan L Marguet, György Buzsáki, and Kenneth D Harris. Sequential structure of neocortical spontaneous activity in vivo. Proceedings of the National Academy of Sciences, 104(1):347–352, 2007.

[22] Shun-ichi Amari. Dynamics of pattern formation in lateral-inhibition type neural fields. Biological cybernetics, 27(2):77–87, 1977.

[23] Hans-Martin R Arnoldi and Wilfried Brauer. Synchronization without oscillatory neurons. Biological Cybernetics, 74(3):209–223, 1996.

[24] John Hertz. Modelling synfire processing. 1997.

[25] Markus Diesmann, Marc-Oliver Gewaltig, and Ad Aertsen. Stable propagation of synchronous spiking in cortical neural networks. Nature, 402(6761):529–533, 1999.

[26] Moshe Abeles, Gaby Hayon, and Daniel Lehmann. Modeling compositionality by dynamic binding of synfire chains. Journal of Computational Neuroscience, 17(2):179–201, 2004.

[27] Arvind Kumar, Stefan Rotter, and Ad Aertsen. Conditions for propagating synchronous spiking and asynchronous firing rates in a cortical network model. Journal of Neuroscience, 28(20):5268–5280, 2008.

[28] Lawrence Christopher York and Mark CW van Rossum. Recurrent networks with short term synaptic depression. Journal of computational neuroscience, 27(3):607, 2009.

[29] Ila R Fiete, Walter Senn, Claude ZH Wang, and Richard HR Hahnloser. Spike-time-dependent plasticity and heterosynaptic competition organize networks to produce long scale-free sequences of neural activity. Neuron, 65(4):563–576, 2010.

[30] Vladimir Itskov, Carina Curto, Eva Pastalkova, and György Buzsáki. Cell assembly sequences arising from spike threshold adaptation keep track of time in the hippocampus. Journal of Neuroscience, 31(8):2828–2834, 2011.

[31] Yao Lu, Yuzuru Sato, and Shun-ichi Amari. Traveling bumps and their collisions in a two-dimensional neural field. Neural Computation, 23(5):1248–1260, 2011.

[32] Amir H Azizi, Laurenz Wiskott, and Sen Cheng. A computational model for preplay in the hippocampus. Frontiers in computational neuroscience, 7:161, 2013.

[33] David Kappel, Bernhard Nessler, and Wolfgang Maass. STDP installs in winner-take-all circuits an online approximation to hidden Markov model learning. PLOS Computational Biology, 10(3), 2014.

[34] Nikolay Chenkov, Henning Sprekeler, and Richard Kempter. Memory replay in balanced recurrent networks. PLoS computational biology, 13(1):e1005359, 2017.

[35] James M Murray et al. Learning multiple variable-speed sequences in striatum via cortical tutoring. Elife, 6:e26084, 2017.

[36] Alexander Seeholzer, Moritz Deger, and Wulfram Gerstner. Stability of working memory in continuous attractor networks under the control of short-term plasticity. PLoS computational biology, 15(4):e1006928, 2019.

[37] Sebastian Spreizer, Ad Aertsen, and Arvind Kumar. From space to time: Spatial inhomogeneities lead to the emergence of spatiotemporal sequences in spiking neuronal networks. PLOS Computational Biology, 15(10):e1007432, October 2019. doi: 10.1371/journal.pcbi.1007432. URL 10.1371/journal.pcbi.1007432.

[38] Carlo Michaelis, Andrew B. Lehr, and Christian Tetzlaff. Robust trajectory generation for robotic control on the neuromorphic research chip loihi. Frontiers in Neurorobotics, 14, November 2020. doi: 10.3389/fnbot.2020.589532. URL 10.3389/fnbot.2020.589532.

[39] Amadeus Maes, Mauricio Barahona, and Claudia Clopath. Learning compositional sequences with multiple time scales through a hierarchical network of spiking neurons. bioRxiv, 2020.

[40] Davide Spalla, Isabel Maria Cornacchia, and Alessandro Treves. Continuous attractors for dynamic memories. eLife, 10, September 2021. doi: 10.7554/elife.69499. URL 10.7554/elife.69499.

[41] Andrew B Lehr, Arvind Kumar, and Christian Tetzlaff. Sparse clustered inhibition projects sequential activity onto unique neural subspaces. bioRxiv, pages 2023–09, 2023.

[42] Rodrigo Perin, Thomas K Berger, and Henry Markram. A synaptic organizing principle for cortical neuronal groups. Proceedings of the National Academy of Sciences, 108(13): 5419–5424, 2011.

[43] X. Jiang, S. Shen, C. R. Cadwell, P. Berens, F. Sinz, A. S. Ecker, S. Patel, and A. S. Tolias. Principles of connectivity among morphologically defined cell types in adult neocortex. Science, 350(6264):aac9462–aac9462, November 2015. doi: 10.1126/science.aac9462. URL 10.1126/science.aac9462.

[44] Xiangmin Xu, Nicholas D. Olivas, Taruna Ikrar, Tao Peng, Todd C. Holmes, Qing Nie, and Yulin Shi. Primary visual cortex shows laminar-specific and balanced circuit organization of excitatory and inhibitory synaptic connectivity. The Journal of Physiology, 594(7):1891–1910, March 2016. doi: 10.1113/jp271891. URL 10.1113/jp271891.

[45] Yangfan Peng, Federico J Barreda Tomas, Paul Pfeiffer, Moritz Drangmeister, Susanne Schreiber, Imre Vida, and Jörg RP Geiger. Spatially structured inhibition defined by polarized parvalbumin interneuron axons promotes head direction tuning. Science Advances, 7(25):eabg4693, 2021.

[46] Simon Weiler, Drago Guggiana Nilo, Tobias Bonhoeffer, Mark Hübener, Tobias Rose, and Volker Scheuss. Orientation and direction tuning align with dendritic morphology and spatial connectivity in mouse visual cortex. Current Biology, 32(8):1743–1753, 2022.

[47] Hemanth Mohan, Matthijs B. Verhoog, Keerthi K. Doreswamy, Guy Eyal, Romy Aardse, Brendan N. Lodder, Natalia A. Goriounova, Boateng Asamoah, A.B. Clementine B. Brakspear, Colin Groot, Sophie van der Sluis, Guilherme Testa-Silva, Joshua Obermayer, Zimbo S.R.M. Boudewijns, Rajeevan T. Narayanan, Johannes C. Baayen, Idan Segev, Huibert D. Mansvelder, and Christiaan P.J. de Kock. Dendritic and axonal architecture of individual pyramidal neurons across layers of adult human neocortex. Cerebral Cortex, 25(12):4839–4853, August 2015. doi: 10.1093/cercor/bhv188. URL 10.1093/cercor/bhv188.

[48] Rajeevan T Narayanan, Robert Egger, Andrew S Johnson, Huibert D Mansvelder, Bert Sakmann, Christiaan PJ De Kock, and Marcel Oberlaender. Beyond columnar organization: cell type-and target layer-specific principles of horizontal axon projection patterns in rat vibrissal cortex. Cerebral cortex, 25(11):4450–4468, 2015.

[49] L Federico Rossi, Kenneth D Harris, and Matteo Carandini. Spatial connectivity matches direction selectivity in visual cortex. Nature, 588(7839):648–652, 2020.

[50] Ye Li, Hui Lu, Pei lin Cheng, Shaoyu Ge, Huatai Xu, Song-Hai Shi, and Yang Dan. Clonally related visual cortical neurons show similar stimulus feature selectivity. Nature, 486(7401):118–121, May 2012. doi: 10.1038/nature11110. URL 10.1038/nature11110.

[51] Zhiwen Ye, Matthew S Bull, Anna Li, Daniel Birman, Tanya L Daigle, Bosiljka Tasic, Hongkui Zeng, and Nicholas A Steinmetz. Brain-wide topographic coordination of traveling spiral waves. bioRxiv, pages 2023–12, 2023.

[52] Walter J Freeman III. Waves, pulses, and the theory of neural masses. Progress in theoretical biology, 2(1):1–10, 1972.

[53] JR Wickens, R Kotter, and ME Alexander. Effects of local connectivity on striatal function: Simulation and analysis of a model. Synapse, 20(4):281–298, 1995.

[54] J Rinzel, D Terman, X-J Wang, and B Ermentrout. Propagating activity patterns in large-scale inhibitory neuronal networks. Science, 279(5355):1351–1355, 1998.

[55] G Bard Ermentrout and David Kleinfeld. Traveling electrical waves in cortex: insights from phase dynamics and speculation on a computational role. Neuron, 29(1):33–44, 2001.

[56] Alex Roxin, Nicolas Brunel, and David Hansel. Role of delays in shaping spatiotemporal dynamics of neuronal activity in large networks. Physical review letters, 94(23):238103, 2005.

[57] Mikhail I Rabinovich, Pablo Varona, Allen I Selverston, and Henry DI Abarbanel. Dynamical principles in neuroscience. Reviews of modern physics, 78(4):1213, 2006.

[58] Doug Rubino, Kay A Robbins, and Nicholas G Hatsopoulos. Propagating waves mediate information transfer in the motor cortex. Nature neuroscience, 9(12):1549–1557, 2006.

[59] Jian-Young Wu, Xiaoying Huang, and Chuan Zhang. Propagating waves of activity in the neocortex: what they are, what they do. The Neuroscientist, 14(5):487–502, 2008.

[60] Evgueniy V Lubenov and Athanassios G Siapas. Hippocampal theta oscillations are travelling waves. Nature, 459(7246):534–539, 2009.

[61] James B Ackman, Timothy J Burbridge, and Michael C Crair. Retinal waves coordinate patterned activity throughout the developing visual system. Nature, 490(7419):219–225, 2012.

[62] Jagdish Patel, Shigeyoshi Fujisawa, Antal Berényi, Sébastien Royer, and György Buzsáki. Traveling theta waves along the entire septotemporal axis of the hippocampus. Neuron, 75 (3):410–417, 2012.

[63] Tatsuo K Sato, Ian Nauhaus, and Matteo Carandini. Traveling waves in visual cortex. Neuron, 75(2):218–229, 2012.

[64] Kazutaka Takahashi, Sanggyun Kim, Todd P Coleman, Kevin A Brown, Aaron J Suminski, Matthew D Best, and Nicholas G Hatsopoulos. Large-scale spatiotemporal spike patterning consistent with wave propagation in motor cortex. Nature communications, 6(1):7169, 2015.

[65] Honghui Zhang and Joshua Jacobs. Traveling theta waves in the human hippocampus. Journal of Neuroscience, 35(36):12477–12487, 2015.

[66] Grégory Faye and Zachary P Kilpatrick. Threshold of front propagation in neural fields: An interface dynamics approach. SIAM Journal on Applied Mathematics, 78(5):2575–2596, 2018.

[67] Randolph F Helfrich, Ian C Fiebelkorn, Sara M Szczepanski, Jack J Lin, Josef Parvizi, Robert T Knight, and Sabine Kastner. Neural mechanisms of sustained attention are rhythmic. Neuron, 99(4):854–865, 2018.

[68] Lyle Muller, Frédéric Chavane, John Reynolds, and Terrence J Sejnowski. Cortical travelling waves: mechanisms and computational principles. Nature Reviews Neuroscience, 19 (5):255–268, 2018.

[69] Zachary W Davis, Lyle Muller, Julio Martinez-Trujillo, Terrence Sejnowski, and John H Reynolds. Spontaneous travelling cortical waves gate perception in behaving primates. Nature, 587(7834):432–436, 2020.

[70] Zachary W Davis, Gabriel B Benigno, Charlee Fletterman, Theo Desbordes, Christopher Steward, Terrence J Sejnowski, John H. Reynolds, and Lyle Muller. Spontaneous traveling waves naturally emerge from horizontal fiber time delays and travel through locally asynchronous-irregular states. Nature Communications, 12(1):6057, 2021.

[71] Sayak Bhattacharya, Scott L Brincat, Mikael Lundqvist, and Earl K Miller. Traveling waves in the prefrontal cortex during working memory. PLoS computational biology, 18(1): e1009827, 2022.

[72] Carlo R Laing, William C Troy, Boris Gutkin, and G Bard Ermentrout. Multiple bumps in a neuronal model of working memory. SIAM Journal on Applied Mathematics, 63(1): 62–97, 2002.

[73] Sebastian Spreizer, Martin Angelhuber, Jyotika Bahuguna, Ad Aertsen, and Arvind Kumar. Activity dynamics and signal representation in a striatal network model with distance-dependent connectivity. Eneuro, 4(4), 2017.

[74] Sen Song, Per Jesper Sjöström, Markus Reigl, Sacha Nelson, and Dmitri B Chklovskii. Highly nonrandom features of synaptic connectivity in local cortical circuits. PLoS Biology, 3(3):e68, March 2005. doi: 10.1371/journal.pbio.0030068. URL 10.1371/journal.pbio.0030068.

[75] Sarah Rieubland, Arnd Roth, and Michael Häusser. Structured connectivity in cerebellar inhibitory networks. Neuron, 81(4):913–929, 2014.

[76] Claudia Espinoza, Segundo Jose Guzman, Xiaomin Zhang, and Peter Jonas. Parvalbumin+ interneurons obey unique connectivity rules and establish a powerful lateral-inhibition microcircuit in dentate gyrus. Nature communications, 9(1):4605, 2018.

[77] Segundo Jose Guzman, Alois Schlögl, Michael Frotscher, and Peter Jonas. Synaptic mechanisms of pattern completion in the hippocampal ca3 network. Science, 353(6304):1117–1123, 2016.

[78] Nicolas Perez-Nieves, Vincent CH Leung, Pier Luigi Dragotti, and Dan FM Goodman. Neural heterogeneity promotes robust learning. Nature communications, 12(1):5791, 2021.

[79] Shira Sardi, Roni Vardi, Yael Tugendhaft, Anton Sheinin, Amir Goldental, and Ido Kanter. Long anisotropic absolute refractory periods with rapid rise times to reliable responsiveness. Physical Review E, 105(1):014401, 2022.

[80] Daniel Miner and Jochen Triesch. Plasticity-driven self-organization under topological constraints accounts for non-random features of cortical synaptic wiring. PLoS computational biology, 12(2):e1004759, 2016.

[81] Kristen A Richardson, Steven J Schiff, and Bruce J Gluckman. Control of traveling waves in the mammalian cortex. Physical review letters, 94(2):028103, 2005.

[82] Leo Hiselius. spreizer-net. https://github.com/leohiselius/spreizer-net, 2021.

[83] Marcel Stimberg, Romain Brette, and Dan FM Goodman. Brian 2, an intuitive and efficient neural simulator. Elife, 8:e47314, 2019.

[84] Carlo Michaelis. Pelenet: a reservoir computing framework for loihi. arXiv preprint 2011.12338, 2020.

[85] Valentino Braitenberg and Almut Schuz. Cortex: Statistics and Geometry of Neuronal Connectivity. Springer Science and Business Media, Germany, 2013. ISBN 3662037335.

[86] Ken Perlin. An image synthesizer. ACM Siggraph Computer Graphics, 19(3):287–296, 1985.

[87] Michael Avermann, Christian Tomm amd Celine Mateo, Wulfram Gerstner, and Carl C. H. Petersen. Microcircuits of excitatory and inhibitory neurons in layer 2/3 of mouse barrel cortex. Journal of Neurophysiology, Volume 107, Article 11, pp. 3116–3134, 2012.

[88] Sandrine Lefort, Christian Tomm, J.-C. Floyd Sarria, and Carl C.H. Petersen. The excitatory neuronal network of the c2 barrel column in mouse primary somatosensory cortex. Neuron, Volume 61, Issue 2, pp 301–316, 2009.

[89] Tomoki Fukai, Vladimir Klinshov, and Jun-Nosuke Teramae. Cortical networks with log-normal synaptic connectivity and their implications in neuronal avalanches. Criticality in Neural Systems, pp. 403–416, 2014.

[90] Carlo Michaelis. Think local, act global: robust and real-time movement encoding in spiking neural networks using neuromorphic hardware. 2022.

[91] Michael Berger, Naubahar Shahryar Agha, and Alexander Gail. Wireless recording from unrestrained monkeys reveals motor goal encoding beyond immediate reach in frontoparietal cortex. Elife, 9:e51322, 2020.

[92] Wulfram Gerstner, Werner M. Kistler, Richard Naud, and Liam Paninski.Neuronal Dynamics: From Single Neurons to Networks and Models of Cognition. Cambridge University Press, USA, 2014. ISBN 1107635195.

